# *In vitro* and *in vivo* CRISPR-Cas9 screens reveal drivers of aging in neural stem cells of the brain

**DOI:** 10.1101/2021.11.23.469762

**Authors:** Tyson J. Ruetz, Chloe M. Kashiwagi, Bhek Morton, Robin W. Yeo, Dena S. Leeman, David W. Morgens, C. Kimberly Tsui, Amy Li, Michael C. Bassik, Anne Brunet

## Abstract

Aging impairs the ability of neural stem cells to transition from quiescence to activation (proliferation) in the adult mammalian brain. Neural stem cell (NSC) functional decline results in decreased production of new neurons and defective regeneration upon injury during aging^1–9^, and this is exacerbated in Alzheimer’s disease^10^. Many genes are upregulated with age in NSCs^3, 11–13^, and the knockout of some of these boosts old NSC activation and rejuvenates aspects of old brain function^14–18^. But systematic functional testing of genes in old NSCs – and more generally in old cells – has not been done. This has been a major limiting factor in identifying the most promising rejuvenation interventions. Here we develop *in vitro* and *in vivo* high-throughput CRISPR-Cas9 screening platforms to systematically uncover gene knockouts that boost NSC activation in old mice. Our genome-wide screening pipeline in primary cultures of young and old NSCs identifies over 300 gene knockouts that specifically restore old NSC activation. Interestingly, the top gene knockouts are involved in glucose import, cilium organization and ribonucleoprotein structures. To determine which gene knockouts have a rejuvenating effect for the aging brain, we establish a scalable CRISPR-Cas9 screening platform *in vivo* in old mice. Of the 50 gene knockouts we tested *in vivo*, 23 boost old NSC activation and production of new neurons in old brains. Notably, the knockout of *Slc2a4*, which encodes for the GLUT4 glucose transporter, is a top rejuvenating intervention for old NSCs. GLUT4 protein expression increases in the stem cell niche during aging, and we show that old NSCs indeed uptake ∼2-fold more glucose than their young counterparts. Transient glucose starvation increases the ability of old NSCs to activate, which is not further improved by knockout of *Slc2a4/*GLUT4. Together, these results indicate that a shift in glucose uptake contributes to the decline in NSC activation with age, but that it can be reversed by genetic or external interventions. Importantly, our work provides scalable platforms to systematically identify genetic interventions that boost old NSC function, including *in vivo* in old brains, with important implications for regenerative and cognitive decline during aging.

## Introduction

The adult mammalian brain contains several neural stem cell (NSC) regions that give rise to newborn neurons and can repair tissue damaged by stroke or brain injuries^1, 2, 9, 19–29^. The most active NSC niche is located in the subventricular zone (SVZ) lining the lateral ventricles of the brain^2, 5, 9, 20, 27, 28, 30–36^. NSCs from the SVZ region can generate thousands of newborn neurons each day in a young adult mouse^20, 31, 33^. The SVZ region comprises a pool of quiescent NSCs that can give rise to activated (proliferating) NSCs, which in turn generate more committed progenitors that migrate out of the niche toward the olfactory bulb, where they differentiate into neurons. The ability of NSCs to activate and form newborn neurons is severely impaired in the aging brain, and this can contribute to deficits in cognition and regeneration^1–8, 30, 37–39^.

Identifying genes that impact NSC activation can lead to interventions that counter tissue defects during aging. Several genetic interventions have been found to improve old NSC activation, including signaling pathways and transcriptional regulators^14–18, 40–44^. However, such studies have been limited in their throughput as they focused on one or a few genes at a time. Thus, we are still missing a systematic understanding of the genes and pathways that functionally affect old NSCs.

More generally, a major challenge in identifying genetic interventions that improve old cells is the establishment of scalable genetic screens in mammals. Aging occurs at both the cell and organismal level, and it is therefore important to develop screens *in vitro* in cells from old organisms and *in vivo* in old tissues. CRISPR-Cas9 genome-wide screens have been developed for a number of phenotypes *in vitro*^45–56^, including with stem cell models of Werner and Hutchinson-Gilford progeria syndrome^57^. However, genetic screens for drivers of aging and rejuvenation in normal old cells have not yet been performed. In addition, *in vivo* genetic screens are challenging in mammals and have been so far limited to cancer^58–65^ and development^66–68^. Thus, developing CRISPR-Cas9 screening platforms for old mammalian cells and organisms has the potential to identify previously unknown gene manipulations that could restore tissue function in older individuals. In the brain, such screens could help identify strategies to counter regenerative and cognitive decline with aging.

## Results

### A genome-wide *in vitro* CRISPR-Cas9 screen identifies over 300 gene knockouts that rejuvenate old NSCs

To systematically identify genes that boost activation of NSCs as a function of age, we developed a genetic screening platform to conduct genome-wide CRISPR/Cas9 knockout screens in primary NSC cultures from young and old mice (Fig. 1a) (see Fig. 2 for counterpart *in vivo* screen). Primary NSC cultures can transition between quiescence (qNSC) and activated (aNSC) states in culture when exposed to different growth factors^69^. Primary qNSC cultures from old mice exhibit decreased ability to activate compared to their young counterparts, recapitulating an *in vivo* aging phenotype^41^. To establish a screening platform, we aged cohorts of mice that express Cas9 (and EGFP) in all cells – hereafter termed ‘Cas9 mice’ (see Methods)^70^. For each independent screen (three total), we harvested NSCs from the subventricular zone (SVZ) of six young (3-4 months) and six old (18-21 months) Cas9 mice. As expected, primary qNSC cultures from old Cas9 mice displayed an impeded ability to activate compared to young counterparts (∼2-fold decline, based on proliferation marker Ki67) (Fig. 1b, c).

**Figure 1.**
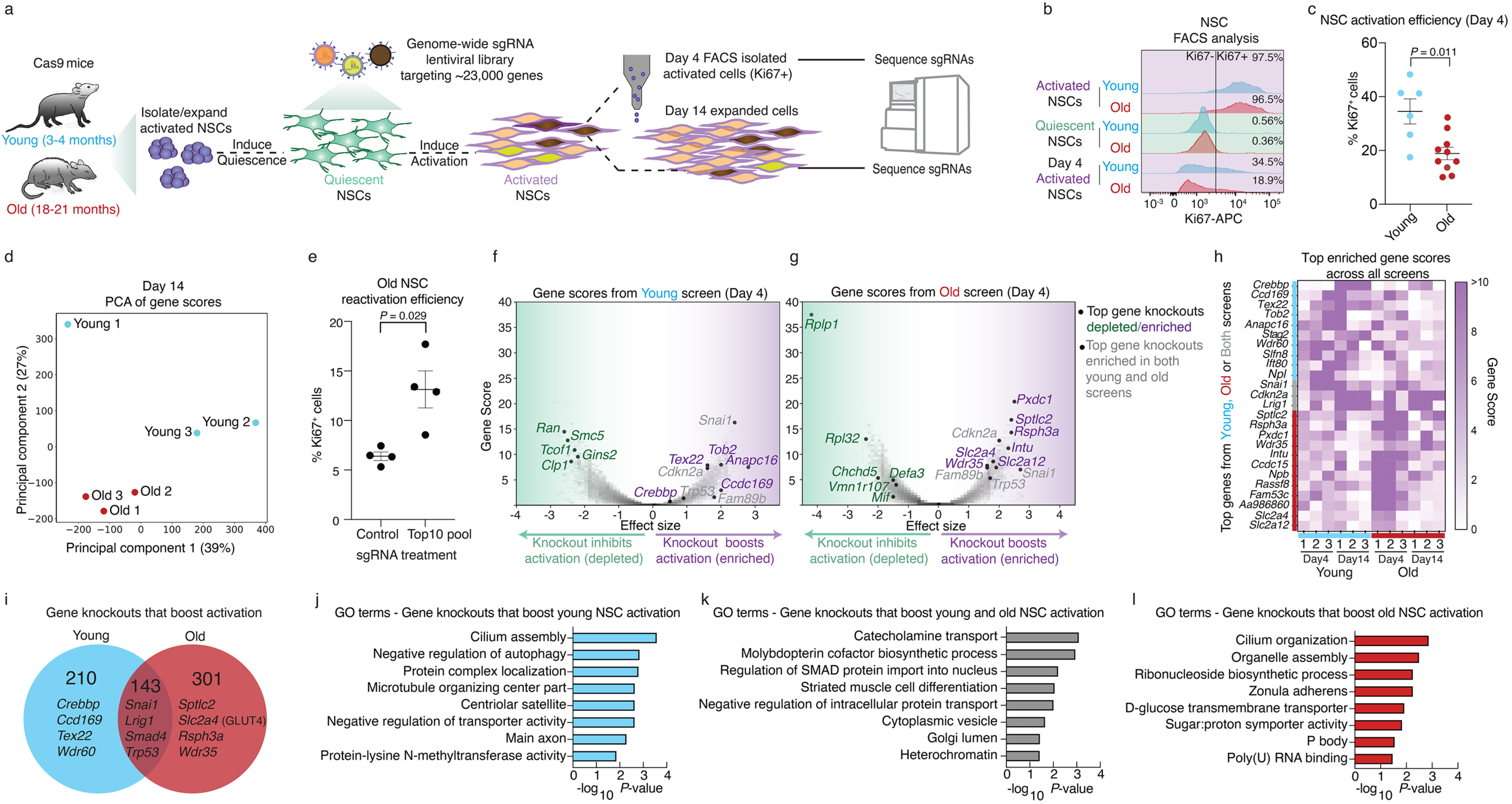
A genome-wide screen identifies 300 gene knockouts that boost old neural stem cell activation. **a,** Screen design overview. Three independent screens were performed with primary NSC cultures from young and old Cas9 mice, each derived from a pool of 6 young (3-4 months old) or 6 old (18-21 months old) Cas9 mice (3 male and 3 female littermates). The active NSC cultures are expanded and then put into a state of quiescence with addition of Bone morphogenetic protein (BMP4) growth factor and removal of EGF growth factor. Quiescent NSCs are infected with a lentiviral genome-wide library of single guide RNAs (sgRNAs) targeting 23,000 genes (with 10 sgRNAs per gene and 15000 control sgRNAs, for a total of 245,000 sgRNAs). Quiescent NSCs are then transitioned from quiescence to activation by removal of BMP4 and addition of EGF growth factor, and then activated NSCs are collected after 4 days (Day 4) by intracellular fluorescence-activated cell sorting (FACS) with a proliferation marker (Ki67) or expanded for 14 days (Day 14). Genomic DNA is prepared and sgRNA constructs are PCR amplified rom Day 4 or Day 14 prior to sequencing on an Illumina Novaseq S4 system. **b,** Example of intracellular FACS analysis with the proliferation marker Ki67 on NSCs from young (3-4 months old, blue) and old (18-21 months old, red) Cas9 mice. Activated NSCs (steady state, top rows), quiescent NSCs (middle rows), or NSCs that transitioned from quiescence to activation (Day 4) that were used for the genome-wide knockout screens (bottom rows). **c,** Quantification of NSC activation efficiency at Day 4 assessed by intracellular Ki67 FACS analysis. Dot plot showing mean +/- SEM of the percentage of cells expressing Ki67. Each dot represents an independent primary NSC culture derived from a pool of six young (3–4 months old) or old (18-21 months old) Cas9 mice (3 male and 3 female littermates). *P*-values determined by two tailed Mann-Whitney test. **d,** Principal Component Analysis (PCA) performed on all gene scores of each of the three independent screens at Day 14: Young 1, 2, 3 (blue) and Old 1, 2, 3 (red). **e**, Quantification of NSC activation efficiency at Day 4 as assessed by FACS with Ki67 (proliferation marker). qNSCs were infected with lentivirus targeting control sgRNAs (100 sgRNAs targeting unannotated genomic regions) or a pool of sgRNAs targeted the top 10 genes (5 sgRNAs per gene) from screens 1 and 2 (FDR <0.1 in both screens. Selecting genes that intersect screen 1 (day 4 or 14) with screen 2 (day 4 or 14)). Each dot represents an independent primary culture of old NSCs derived from a pool of six old (18-21 months old) Cas9 mice (3 male and 3 female littermates). *P*-values determined by two tailed Mann-Whitney test. **f, g** Volcano plots of example screen results (screen 2) at Day 4 for young **(f)** or old NSCs **(g)**, showing gene scores (y-axis) in relation to effect size (x-axis). Each dot represents one gene. Labelled dots are top ranking genes (FDR<0.1 in at least 2 of the 3 independent screens, across any time point (day 4 or day 14)) whose knockouts boost NSC activation (purple, corresponding to enriched sgRNAs) or impede NSC activation (green, corresponding to depleted sgRNAs) in an age-dependent manner or whose knockouts boost activation regardless of age (grey, corresponding to enriched sgRNAs). See Supplementary Table 1 for complete list of gene scores. **h,** Heatmap showing gene scores across samples and days for the top 10 gene knockouts that boost young or old activation, with addition of *Slc2a4* (GLUT4), *Slc2a12* (GLUT12), and 3 gene knockouts that boost both young and old activation (*Snai1*, *Cdkn2a*, *Lrig1*), as identified across all screens. **i,** Venn diagram of all gene knockouts that boost NSC activation in at least 2 of the 3 independent screens (FDR<0.1) in young (blue) or old NSCs (red). **j-l)** Selected Gene Ontology (GO) terms associated with the genes knockouts that boost young (**j**), young and old (**k**) or old (**l**) NSC activation (FDR<0.1 in at least 2 of the 3 independent screens), assessed using EnrichR, focusing on the GO “cell component”, “molecular function”, and “biological process” libraries. Complete lists of GO terms in Supplementary Table 2. *P-*values calculated by EnrichR, using a Fisher’s exact test.

**Figure 2.**
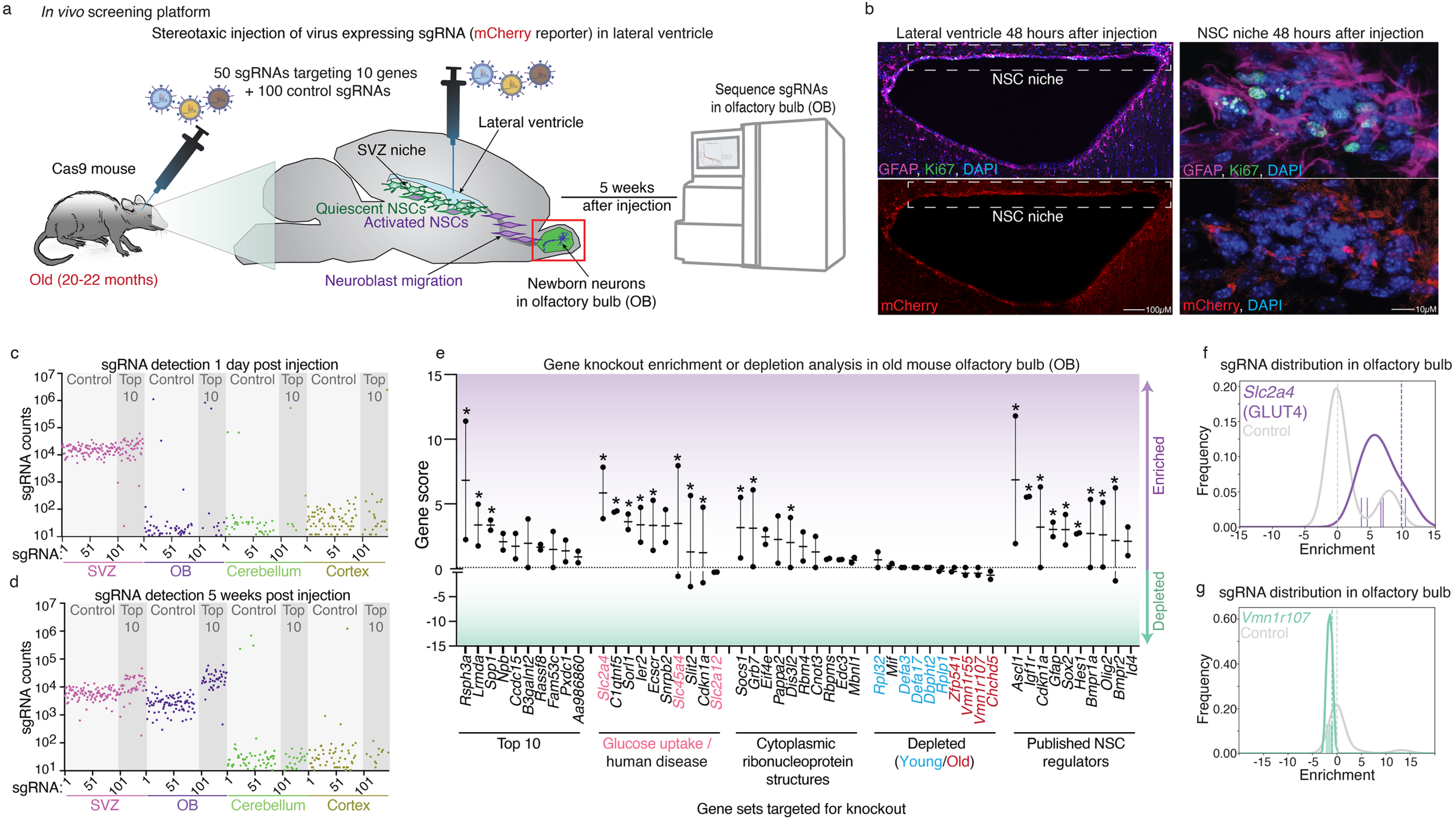
*In vivo* screening platform for rapid testing of gene knockout effects on neural stem cell activation. **a,** Overview of *in vivo* screen platform design in old mice. Old (∼20-22 months old) Cas9 mice (all female) are injected with lentivirus expressing sgRNAs directly into the lateral ventricles (in close proximity to the SVZ niche). In each case, the sgRNAs target 10 genes of interest (5 sgRNAs per gene) along with 100 control sgRNAs (targeting unannotated genomic regions), for a total of 150 sgRNAs (pool of lentivirus that each express one of the 150 sgRNAs). 5 weeks after injection, genomic DNA is harvested from the olfactory bulb and sgRNAs are sequenced to determine enrichment or depletion. **b,** Immunofluorescence images of sections of the subventricular zone (SVZ) neural stem cell niche of old (21-months old) Cas9 mouse brain, 48 hours after stereotactic injection of lentiviruses expressing mCherry and 100 control sgRNAs (pool of lentivirus that each express one of the 100 control sgRNAs). Markers: mCherry (lentivirus-infected cells, red), GFAP (NSC and astrocyte marker, magenta), Ki67 (proliferative cell marker, green), and DAPI (nuclei, blue). Right panels: high magnification images of SVZ NSC niche infected with virus. **c-d,** Normalized sgRNA counts of the Top 10 gene pool (50 sgRNAs) and control (100 sgRNAs) libraries in various brain regions, either 1 day (**c**) or 5 weeks (**d**) post virus injection. The Top 10 gene pool corresponds to genes that when knocked-out boosted old NSC activation in our *in vitro* screens (screen 1 and 2). All sgRNA counts (measured by high throughput sequencing) are normalized to total counts between brain regions to account for variance in sequencing depth. **e,** CasTLE gene scores for 50 gene knockouts tested in groups of 10 in old (20-22 months old) female Cas9 mice olfactory bulbs. Each sgRNA library of 10 genes was injected into 3 old Cas9 mice, one mouse was used for the 1 day post injection SVZ sequencing of starting sgRNA pool, and the other two mice were left for 5 weeks and then the olfactory bulb sgRNAs were sequenced. Olfactory bulb sgRNA enrichment was computed with CasTLE by comparison to the 24-hour SVZ sequenced sample. Dot plot showing mean +/- SEM of CasTLE score in two independent mice. Each dot represents gene score from one mouse. *: gene hits with a 95% confidence interval that did not contain 0 as computed by CasTLE analysis. **f, g**, Histograms plotting the relative enrichment and frequency of each sgRNA targeting *Slc2a4* (GLUT4, purple) *or Vmn1r107* (green) or the control (grey) sgRNA pool. Hashed line indicates the CasTLE computed relative enrichment effect size for the sgRNA targeting the gene of interest relative to control sgRNA pool. Colored dashes above the x-axis represent each of the 5 sgRNA targeting *Slc2a4* (GLUT4) *or Vmn1r107* and their relative enrichment. See also Supplementary Table 3.

We performed three independent genome-wide CRISPR-Cas9 screens for genes that impact qNSC activation as a function of age. To this end, we expanded NSCs from young and old Cas9 mice, induced quiescence, and then transduced over 400 million qNSCs with lentiviruses that express a single guide RNA (sgRNA) library targeting all ∼23,000 protein coding genes in the genome, with 10 unique sgRNAs per gene, as well as 15,000 control sgRNAs (∼245,000 sgRNAs total)^46^ (Fig. 1a). Five days after sgRNA library transduction, qNSCs were activated with growth factors and expanded for either 4 or 14 days and processed as independent samples for sequencing (Fig. 1a). The day 4 time point was chosen to isolate any cell that successfully activated by fluorescence-activated cell sorting (FACS) based on the proliferation marker Ki67. The day 14 timepoint allowed for more robust enrichment of cells with knockouts that maintained self-renewal capabilities over long periods of time, outgrew other cells, and therefore did not require FACS isolation. After 4 or 14 days, we generated libraries of sgRNAs from all cells and processed them for high-throughput sequencing. The sgRNA libraries had sufficient coverage of the sgRNA pool for genome-wide screen analysis (Extended Data Fig. 1a, b).

To analyze each screen sample, we assessed sgRNA enrichment/depletion by CasTLE analysis, which uses the sequencing read counts of the 10 sgRNAs targeting each gene and compares to the control sgRNA distributions to compute effect size, gene score, confidence interval, and *P*-value^45^. Interestingly, principal component analysis (PCA) on CasTLE gene score values separated our three independent screens based on the age of NSCs (Fig. 1d and Extended Data Fig. 1c, d), highlighting aging as a primary contributor to the screen’s outcome. We independently verified that a sub-screen of a pool of the top 10 gene knockouts that enriched in old NSC activation in screens 1 and 2 indeed boosted NSC activation as assessed by Ki67^+^ FACS analysis (Fig. 1e). We then directly compared young and old NSC screens to identify the gene knockouts that boost NSC activation specifically in young NSCs, specifically in old NSCs, or regardless of age (Fig. 1f-i, Extended Data Fig. 1e, f and Supplementary Table 1).

Overall, our genome-wide screens identified 654 and 1,386 genes whose knockouts boosted or impeded NSC activation respectively (FDR<0.1 in 2 or more independent screens, Supplementary Table 1). We found 143 gene knockouts that enhanced both young and old NSC activation, 210 specific to young NSC activation, and 301 specific to old NSC activation (Fig. 1i and Supplementary Table 1). Consistent with previous findings, genes implicated in the maintenance of NSC quiescence (e.g. *Snai1, Smad4* (TGFß pathway), *and Lrig1*)^71–73^ as well as general cell cycle regulators (e.g. *Trp53*, *Cdkn2a*)^74, 75^ were enriched as knockouts that increase both young and old NSC activation (Fig. 1h, i and Supplementary Table 1). Surprisingly, a large fraction of gene knockouts increased activation only in young or only in old NSCs, suggesting some NSC quiescence regulators change as a function of age. Many of the old specific gene knockouts had not been previously implicated in NSC activation, such as the genes encoding *Sptlc2*, *Rsph3a*, *Pxdc1* and the glucose transporters *Slc2a4* (GLUT4) and *Slc2a12* (GLUT12) (Fig. 1f-i and Extended Data Fig. 2a-c).

GO term analysis of gene knockouts that specifically boost old NSC activation revealed cilium organization, cytoplasmic ribonucleoprotein structures (P bodies, RNA binding proteins), and glucose transport (Fig. 1l and Supplementary Table 2. See Extended Data Fig. 2 d-g for genes that impede NSC activation). Primary cilia are linked to the quiescent state of NSCs^76^ and knockout of primary cilia genes in young NSCs leads to decreased proliferation and maintenance in mice^77–,81^, but their role in aging NSCs is not known. The cytoplasmic ribonucleoprotein structures GO term is interesting with regard to the cytoplasmic protein aggregate structures that are observed in aging quiescent NSCs^41^. Glucose metabolism signatures have been linked to the quiescent state of NSCs^23, 41, 82, 83^ and regulation of glucose metabolism has been shown to impact NSC self-renewal and differentiation to neurons^83–87^, but glucose transport and metabolism have not been previously implicated in old qNSC activation. Together, these results provide an exhaustive dataset with gene knockouts that restore the transition from quiescence to activation, a key aspect of neural stem cell aging, and it includes many previously unknown genes and pathways.

### Development of an *in vivo* CRISPR-Cas9 screening platform to rapidly screen gene knockouts for their ability to rejuvenate NSC activity in the old brain

Primary cultures from old mice recapitulate several aspects of *in vivo* aging but not all, and there are no current methods to rapidly screen multiple genes for functional impact on aging cells *in vivo.* To test if gene knockouts could also boost NSC function *in vivo*, we developed a gene knockout screening platform for the aging brain. The SVZ neurogenic region provides a good paradigm for *in vivo* screening. Quiescent NSCs of the SVZ normally activate and produce progeny in this niche, which then migrate out of the niche to generate new neurons at a distal location in the olfactory bulb^9^. We leveraged the natural properties of this regenerative region to design a CRISPR-Cas9 screening platform in old mice. We performed stereotaxic brain surgery in old Cas9 mice to inject sgRNA- and mCherry-expressing lentiviruses directly into the lateral ventricles, in close proximity to NSCs from the SVZ neurogenic niche (Fig. 2a). We verified that injection led to infection of NSCs in the niche by mCherry immunofluorescence and staining for markers of quiescent NSCs (GFAP^+^, Ki67^-^) and activated NSCs (GFAP^+^, Ki67^+^) (Fig. 2b and Extended Data Fig. 3a). After 5 weeks to allow time for both knockout to occur and NSC progeny to migrate to the olfactory bulb, we collected the entire olfactory bulb for sequencing of sgRNAs (Fig. 2a). We verified that injecting sgRNAs targeting EGFP in the SVZ niche of old Cas9 mice (which also express an EGFP reporter) led to decreased EGFP staining in the olfactory bulb after 5 weeks (Extended Data Fig. 3b, c).

We verified coverage and diversity of sgRNAs *in vivo* by injecting old Cas9 mice with lentivirus expressing 50 sgRNAs targeting the top 10 genes (‘Top 10’ library) that specifically boosted old NSC activation from our first two *in vitro* screens (each gene targeted by 5 unique sgRNAs), and 100 negative control sgRNAs targeting unannotated regions of the genome (150 sgRNAs total). One day after injection, we extracted and sequenced genomic DNA from the SVZ niche (and 3 other distant brain regions as a control) (Fig. 2c). At the day one time point, 150 sgRNAs were detected in the SVZ niche and only a fraction of sgRNAs at 100-1000 fold fewer sequencing reads were detected in the olfactory bulb, cerebellum, and outer cortex (Fig. 2c and Extended Data Fig. 4a). Five weeks after injection (in a different old Cas9 mouse), we also collected the same brain regions (Fig. 2d). At the week five time point, 150 sgRNAs were again detected in the SVZ niche, but now the majority were also detected in the olfactory bulb, suggesting NSCs containing sgRNAs had migrated out of the SVZ niche and allowed distal neurogenesis in the olfactory bulb (Fig. 2d and Extended Data Fig. 4b). Importantly, the 50 sgRNAs targeting the Top 10 gene pool were strongly enriched over the 100 control sgRNAs in the olfactory bulb, and to a lesser extent in the SVZ niche (Fig. 2c, d). The sgRNA abundance in the olfactory bulb is unlikely to be explained by expansion of cells directly infected in the olfactory bulb itself given the lack of sgRNA diversity in the olfactory bulb (OB) at one day post-infection (Fig. 2c and Extended Data Fig. 4a). Importantly, we also did not observe Top 10 sgRNA pool enrichment over control in the cerebellum or cortex (Fig. 2d) or in wildtype mice that do not express Cas9 (Extended Data Fig. 4c). Together, these data indicate that our *in vivo* platform leverages the natural properties of the NSC niche and the production of neurons in a distal region, which allows efficient targeted CRISPR-Cas9 screening *in vivo*.

Using this platform, we performed 5 knockout screens in old mice, targeting groups of 10 genes for their ability to boost NSC activity and migration to the olfactory bulb. In addition to the Top 10 gene library, we selected 4 other sets of 10 genes based on our genome-wide screens in cultured NSCs. The 5 libraries were: i) ‘Top 10’ library, with genes that when knocked-out boosted old NSC activation in our *in vitro* screens (screen 1 and 2) (e.g. *Rsph3a*), ii) ‘Glucose uptake/human disease’ library, with genes that when knocked-out also boosted activation of old NSCs in our *in vitro* screens (e.g. *Slc2a4* (GLUT4), *Slc2a12* (GLUT12), *Sorl1*), iii) ‘Cytoplasmic ribonucleoprotein structures’ library, with genes that belong to the P bodies and cytoplasmic ribonucleoprotein/stress granule GO terms, some of which boosted old NSC activation in our *in vitro* screen (e.g. *Dis3l2, Mbnl1, Edc3*), iv) ‘Depleted (Young/Old)’ library, with genes that when knocked-out blocked young or old NSC activation in our *in vitro* screens (screen 1 and 2), and v) ‘Published NSC regulators’ library, with genes that have previously been implicated in NSC function in the literature (Fig. 2e). We injected old Cas9 mice with virus to express one of the 5 libraries along with the 100 control sgRNA library, waited five weeks, and then sequenced sgRNAs in the olfactory bulbs and performed CasTLE analysis (Fig. 2e). Of the 50 gene knockouts tested, we identified 23 gene knockouts that were significantly enriched in the olfactory bulb, suggesting that they boost old NSC activation (and/or migration and differentiation) (e.g., *Rsph3a, Slc2a4, Socs1*) (Fig. 2e, f and Extended Data Fig. 4 d, e, g). Of the 10 gene knockouts predicted to impede NSCs activation, the 4 predicted to specifically block old NSC activation were slightly depleted in the olfactory bulb but did not reach significance (Fig. 2e, g and Extended Data Fig. 4f). Interestingly, of the activating gene knockouts tested based on our *in vitro* screens, some of the most abundant and enriched sgRNAs targeted genes associated with cilia (*Rsph3a*), glucose uptake (*Slc2a4, Slc45a4*), and Alzheimer’s disease (*Sorl1*) (Fig. 2e). The ‘Glucose uptake/human disease’ library was most strongly enriched, with 8 of the 10 gene knockouts reaching significance (Fig. 2e). Overall, these data establish a scalable platform to genetically screen *in vivo* for genes that impact old NSC function and highlight the importance of cilia, glucose metabolism, and ribonucleoprotein structures for aging NSCs.

### Knocking-out the glucose transporter GLUT4, which increases with age, boosts neurogenesis in old mice

One of the most consistent gene knockouts from our *in vivo* screens that boost old NSC function both *in vitro* and *in vivo* is *Slc2a4* (GLUT4), which encodes an insulin-dependent glucose transporter protein^88^. While glycolysis and glucose uptake regulate NSC proliferation and differentiation to neurons^84–86^, the importance of glucose uptake – and the GLUT4 transporter – in aged NSC function has never been examined. We asked if reducing GLUT4 in the SVZ niche is sufficient to boost the ability of NSCs to generate newborn neurons in the olfactory bulb (neurogenesis) in old individuals. To this end, we stereotactically injected sgRNAs targeting *Slc2a4* (GLUT4), *Vmn1r107* (a gene that was depleted in our *in vitro* and *in vivo* NSC activation screens), or unannotated regions of the genome (control), into the ventricles of old Cas9 mice. We verified that sgRNAs targeting *Slc2a4* (GLUT4) successfully reduced GLUT4 staining in the olfactory bulb five weeks after injection of lentivirus in the SVZ NSC niche (Extended Data Fig. 5a, b). To track newborn cells arriving in the olfactory bulb, we injected mice weekly starting 7 days post virus injection with EdU (5-ethynyl-2’-deoxyuridine), a thymidine analog that incorporates into newly synthesized DNA and can be visualized by immunofluorescent assays (Fig. 3a). We first quantified the number of newborn cells in the olfactory bulb five weeks after virus injection by co-staining for EdU (newborn cells) and mCherry (sgRNA lentivirus reporter) (Fig. 3b). *Slc2a4* (GLUT4) knockout resulted in over 2-fold increase in the proportion of newborn cells in the olfactory bulb that were mCherry^+^ relative to control treatment (Fig. 3b). We then assessed how well cells with GLUT4 knockout differentiated into neurons by staining for NeuN, a marker of mature neuron nuclei^89^. The majority of cells targeted by the *Slc2a4* (GLUT4) sgRNA (mCherry^+^) in the olfactory bulb were positive for NeuN (Fig. 3c) with no significant change in NeuN expression levels or proportion of newborn (EdU^+^) cells that were NeuN^+^, relative to control treatment (Fig. 3d and Extended Data Fig. 5c). Collectively, these results suggest that *Slc2a4* (GLUT4) knockout in the niche improves neurogenesis.

**Figure 3.**
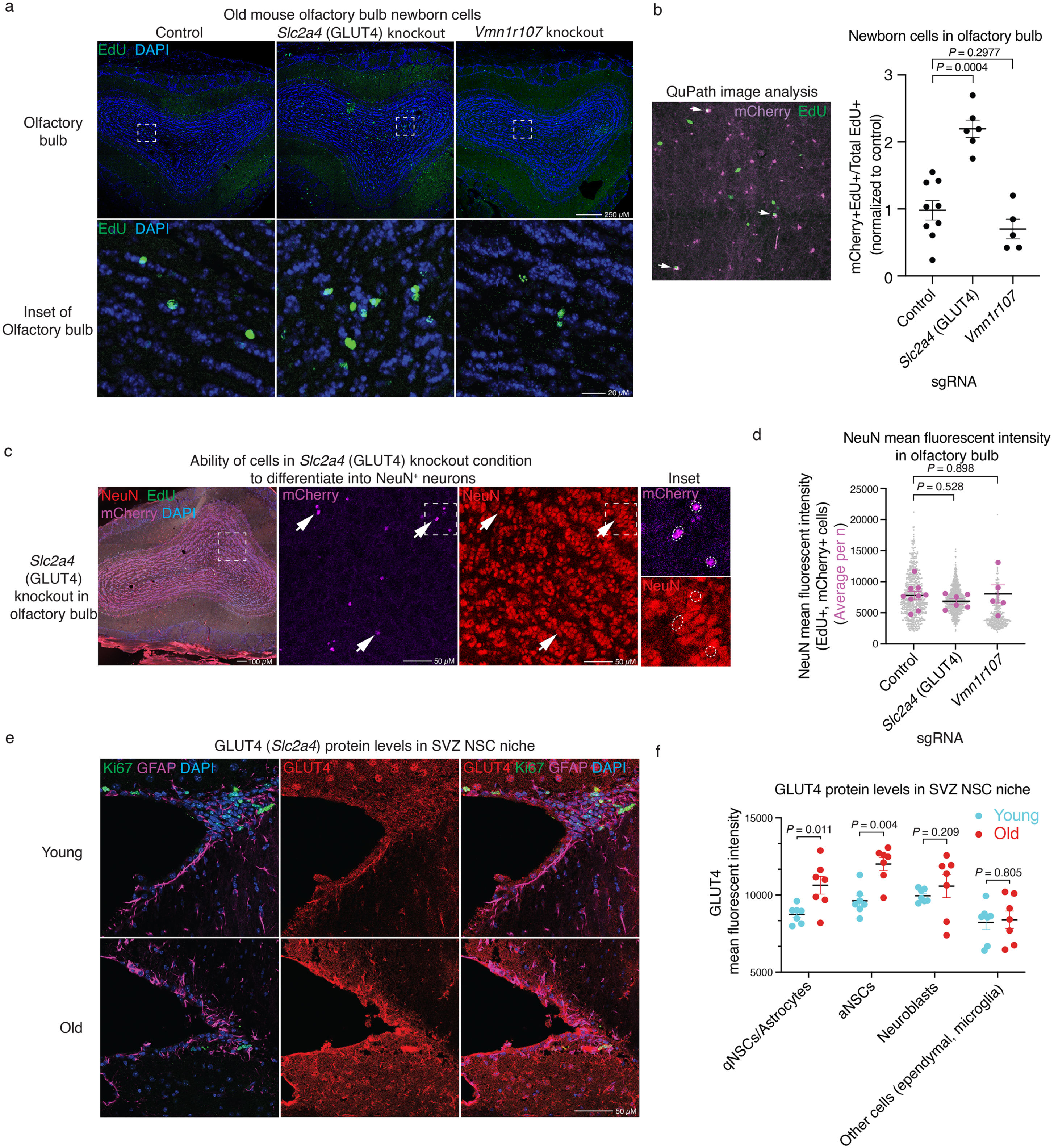
*Slc2a4* (GLUT4) knockout in the SVZ neural stem cell niche boosts neurogenesis in old mice. **a,** Representative immunofluorescence images of olfactory bulb sections from old (∼20 months old) male Cas9 mice, 5 weeks after injection of lentivirus expressing control (unannotated genomic regions), *Slc2a4* (GLUT4) or *Vmn1r107* sgRNAs directly into the lateral ventricles. Mice were injected with EdU once per week, starting one week after virus injection, for 4 weeks. Markers: EdU (newborn cells, green) and nuclei (DAPI, blue). Zoomed-out image with the dashed white square representing the inset (top) and zoomed-in view as an inset (bottom). **b,** Example image of mCherry (lentivirus-infected cells, magenta) and EdU (newborn cells, green) staining (left) and QuPath image quantification of the average number of EdU^+^ and mCherry^+^ cells in the olfactory bulbs, normalized to total EdU^+^ cells (right) in old Cas9 mice (18-23 months old) (for ages and sex, see Supplemental Data 4). Dot plot, with each dot representing counts from one mouse, showing mean +/- SEM. Results are from 3 independent experiments for a total of n = 5-9 mice. Each dot represents the average number of cell counts for one mouse, from 3 serial sections taken at 100 µm intervals across the olfactory bulb pair, with the treated conditions normalized to average of control. *P*-values determined by two-tailed Mann-Whitney test. **c,** Representative immunofluorescence images of olfactory bulb sections from old (20 months old) male Cas9 mouse, 5 weeks after injection of lentivirus expressing *Slc2a4* (GLUT4) sgRNAs directly into the lateral ventricles. Mice were injected with EdU once per week, starting one week after virus injection, for 4 weeks. Markers: EdU (newborn cells green), mCherry (lentivirus-infected cells, magenta), NeuN (mature neuron marker, red), and DAPI (nuclei, blue). White arrows highlight NeuN^+^ cells that are infected with lentivirus (expressing one of the sgRNAs targeting GLUT4). **d,** QuPath image quantification of NeuN mean fluorescent intensity in infected (mCherry^+^) newborn cells (EdU^+^) in the olfactory bulb of old Cas9 mice (18-23 months old) (mix of males and females, see Extended Data *raw data* file), 5 weeks after injection of lentivirus expressing control (unannotated genomic regions), *Slc2a4* (GLUT4), or *Vmn1r107* sgRNAs directly into the lateral ventricles. Mice were injected with EdU once per week, starting one week after virus injection, for 4 weeks. Dot plot showing mean +/- SEM of quantification results from 3 independent experiments, each with ∼2-3 mice, for a total of n = 5-9 mice. Each lavender dot represents the average NeuN fluorescence intensity of all enumerated cells in one mouse, from 3 serial sections taken at 100 µm intervals across the olfactory bulb pair. Each grey dot represents a single cell NeuN fluorescence intensity, showing all cells across all samples for each respective treatment. Average NeuN fluorescence intensity of cells (lavender dots) of each treated mouse was statistically compared between treated and control mice. *P*-values determined by two tailed Mann-Whitney test. **e,** Representative immunofluorescence images of coronal sections of the SVZ neural stem cell niche from young (3-4 months old) and old (18-21 months old) male Cas9 mice. Markers: GLUT4 (red), Ki67 (proliferation maker, green), GFAP (NSC and astrocyte marker, magenta), and DAPI (nuclei, blue). Cell types were identified as follows: qNSC/astrocyte (GFAP^+^/Ki67^-^), aNSC (GFAP^+^/Ki67^+^), Neuroblast (GFAP^-^/Ki67^+^) and other cells (ependymal, microglia; GFAP^-^/Ki67^-^). **f,** QuPath image quantification of GLUT4 mean fluorescence intensity in cells of the neural stem cell niche from 7 young (3-4 months old) and 7 old (18-21 months old) male Cas9 mice. Cell types were identified as in e. Dot plot showing mean +/- SEM of quantification results from 2 independent experiments, each with 3-4 mice, for a total of n = 7 mice. Each dot represents GLUT4 mean fluorescent intensity average of cells from one mouse. *P*-values determined by two tailed Mann-Whitney test.

Given the beneficial effect of GLUT4 knockout in old NSCs *in vivo*, we asked whether GLUT4 protein expression itself changes in the SVZ neurogenic niche with age. Immunofluorescence staining of SVZ sections from young and old mice revealed that GLUT4 protein increased both in quiescent NSCs/astrocytes (GFAP^+^Ki67^-^) and activated NSCs/NPCs (GFAP^+^Ki67^+^) in the old brain (Fig. 3e, f and Extended Data Fig. 5d). In contrast, GLUT4 protein expression did not change with age in other cell types (ependymal, microglia; GFAP^-^Ki67^-^) (Fig. 3e, f and Extended Data Fig. 5d (control)). Thus, the glucose transporter GLUT4 increases in expression during aging in NSCs *in vivo*, and this could explain at least in part the decline in neurogenesis in old brains.

### Old NSCs exhibit high glucose uptake which can be targeted to ameliorate activation

GLUT4 (*Slc2a4*) is a transporter that increases glucose uptake in an insulin-dependent manner in cells^88^ (Fig. 4a). We therefore assessed components of the glucose uptake pathway (insulin-dependent or independent) in the context of NSC aging by mining our genome-wide *in vitro* screen results. Among the 12 known glucose transporters, only *Slc2a4* (GLUT4) and *Slc2a12* (GLUT12), when knocked out *in vitro* (and *in vivo* for GLUT4), significantly boosted old NSC activation (Fig. 4a, b). Moreover, *Stx4a*, which encodes a protein that facilitates the fusion of GLUT4 storage vesicles with the plasma membrane^88^, boosted old NSC activation in one of the screens (Fig. 4b). Consistently, GLUT4 and STX4A proteins were expressed at higher levels in old quiescent NSCs in culture (Fig. 4c-e and Extended Data Fig. 6a), and *in vivo* (see Fig. 3e, f for GLUT4 and Extended Data Fig. 6b, c for STX4A). In contrast, proteins in the glycolysis, insulin and associated downstream pathways (e.g. hexokinase enzymes HK1-3, mTOR, GSK3B and FOXO) were not enriched in the genome-wide screens in old NSCs, and some insulin pathway genes were even depleted (Fig. 4a, b). Together, these observations suggest that glucose transporter activity, rather than upstream signaling, plays a central role in regulating old NSC activation.

**Figure 4.**
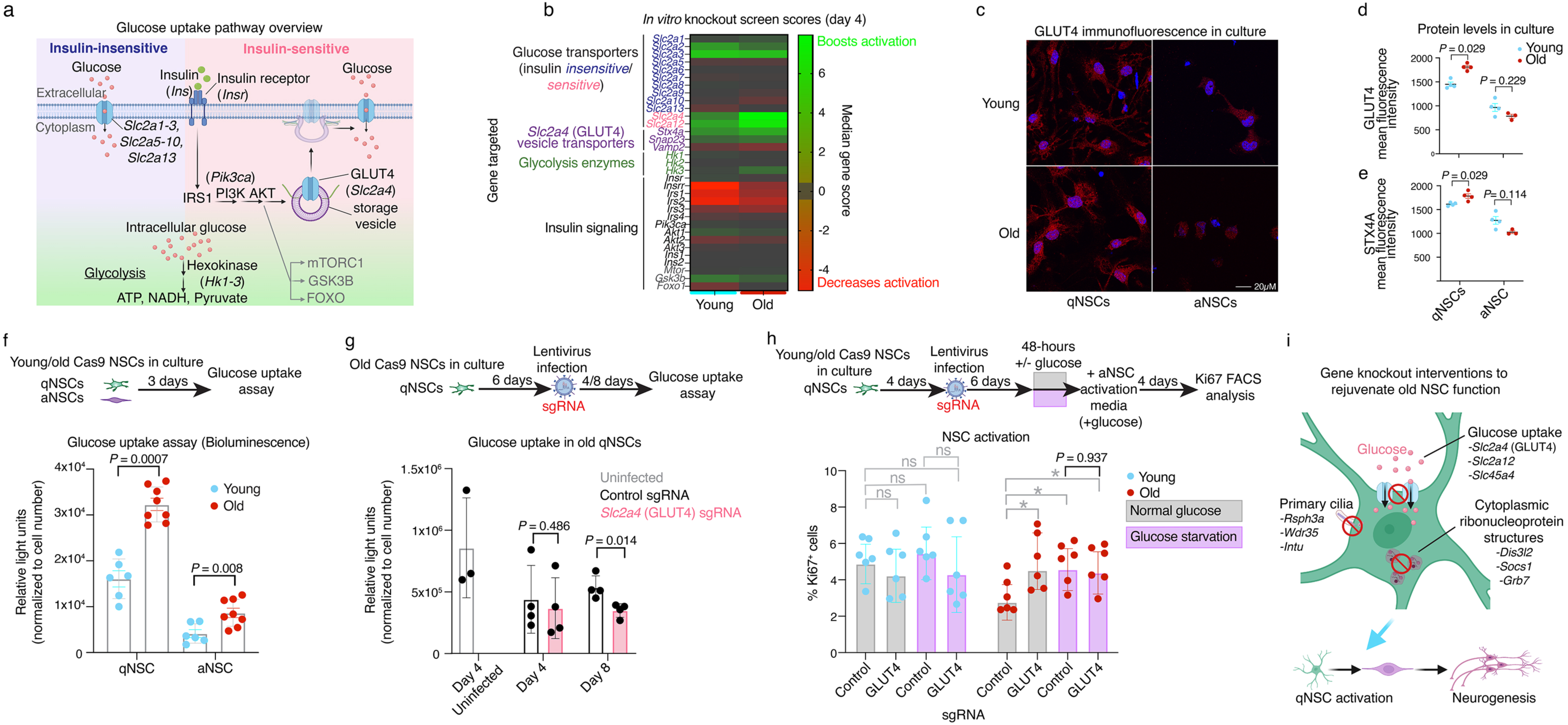
Old NSCs exhibit high glucose uptake which can be targeted to ameliorate activation. **a,** Overview of the glucose uptake pathway. **b,** Heatmap plotting the glucose and insulin pathway median gene scores (CasTLE) from our three genome-wide *in vitro* screens at the Day 4 time point. Gene knockouts that boost (green) or impede (red) activation. **c-e,** Example immunofluorescence image for GLUT4 (**c**) and quantification of primary NSC cultures from young (3-4 months old) or old (18-21 months old) Cas9 mice (mixed sex), either plated in quiescence NSC media (qNSCs) for 7 days prior to imaging or plated in activation NSC media (aNSCs) 2 days prior to imaging. Markers: GLUT4 (**c**, **d,** red), STX4A (Extended Data Fig. 6a, **e,** green) and DAPI (nuclei, blue). QuPath image quantification of GLUT4 or STX4A mean fluorescence. Dot plot showing mean +/- SEM of results from 3-4 independent NSC cultures derived from a pool of 6 mixed-sex mice. Each dot represents an independent NSC culture. *P*-values determined by two-tailed Mann-Whitney test. **f,** Bioluminescent glucose uptake assay on primary qNSCs or aNSCs from young (3-4 months old) or old (18-21 months old) mice. Dot plot showing mean +/- SEM of results from 6-8 independent NSC cultures, each from a pool of 6 mice, 3 males and 3 females. Each dot represents an independent NSC culture. *P*-values determined by two-tailed Mann-Whitney test. **g,** Bioluminescent glucose uptake assay on primary qNSC cultures from old (18-21 months old) mice treated with lentivirus expressing *Slc2a4* (GLUT4) or control (unannotated genomic regions) targeting sgRNAs, or uninfected (no treatment, NT). NSCs were plated in qNSC media 6 days prior to infection with lentivirus to express sgRNA and assessed for glucose uptake either at Day 4 or Day 8 after infection. Dot plot showing mean +/- SEM of results from 3-4 independent NSC cultures, each from a pool of 6 mice (3 males and 3 females). Each dot represents an independent NSC culture. *P*-values determined by two-tailed Mann-Whitney test. **h**, Percentage of qNSCs from mixed sex young (3-4 months old) or old (18-21 months old) that successfully activated (% Ki67^+^) 4 days after transition to aNSC media, as assessed by Ki67 intracellular FACS analysis. NSCs were placed in qNSC media for 4 days, exposed to lentivirus to express sgRNAs targeting *Slc2a4* (GLUT4) or unannotated genomic regions (control). Then 6 days later the cell media was replaced with qNSC media with (grey) or without (pink) glucose for 48 hours, at which point the media was replaced with aNSC media with glucose and the cells were allowed to activate for 4 days prior to intracellular FACS analysis with Ki67. Dot plot showing mean +/- SEM of results from 6 independent NSC cultures, each from a pool of 6 mice, 3 males and 3 females. Each dot represents an independent NSC culture. *P*-values determined by two-tailed Mann-Whitney test. **i**, Summary figure of gene knockouts, and their associated cellular processes, that boost old NSC activation.

We asked if glucose uptake changes with age in NSCs and if this can be manipulated to improve old NSC activation. Glucose uptake assays showed that both old quiescent and active NSCs in culture uptake ∼2-fold more glucose than their young counterparts (Fig. 4f and Extended Data Fig. 6d). Quiescent NSCs in general had higher levels of glucose uptake than activated NSCs (Fig. 4f and Extended Data Fig. 6d), suggesting less reliance on glucose as NSCs transition from quiescence to activation. We therefore asked if restricting glucose uptake, either genetically or by depleting glucose in the media, benefits the activation of old NSCs. *Slc2a4* (GLUT4) knock-out reduced glucose uptake in old quiescent NSCs (Fig. 4g and Extended Data Fig. 6e) and improved the ability of old quiescent NSCs to transition from quiescence to activation (Fig. 4h). Moreover, a brief 48-hour pulse of glucose starvation in quiescent NSCs improved their ability to activate out of quiescence (Fig. 4h and Extended Data Fig. 6f). Importantly, GLUT4 knockout did not further increase old qNSC activation in the context of a pulse of glucose starvation (Fig. 4h and Extended Data Fig. 6f), suggesting that one mechanism by which GLUT4 reduction improves old NSC activation is by lowering glucose. Hence, old quiescent NSCs are in a state of high glucose uptake likely due to increased GLUT4/STX4A expression during aging, but this can be reversed through genetic or glucose restriction perturbations to ameliorate old qNSC activation. These results validate the use of genetic screens in old cells to uncover genetic pathways (e.g. glucose metabolism, primary cilia, ribonucleoprotein structures) and interventions to boost NSC function during aging (Fig. 4i).

## Discussion

This study represents the first genome-wide knockout screen to identify genes that impact aging and rejuvenation in a regenerative neurogenic niche. It also shows the development of a new CRISPR-Cas9 targeted screening platform to study functional drivers of cellular aging *in vivo*. The power of knockout screens is their ability to uncover new biology. Here we uncover over 300 genes that when knocked out boost the activation of old NSCs *in vitro*, including glucose metabolism, ribonucleoprotein structures, and primary cilia associated genes as well as many other genes. These genes contribute to the mechanisms allowing NSCs to transition out of quiescence into activation – a step that is defective in aging (Fig. 4i). Most of these genes had not previously been identified to regulate NSCs, especially in old cells, and they represent a resource for functional regulators of NSC aging and rejuvenation.

The knockout of insulin-sensitive glucose transporter GLUT4 was consistently a top hit for both *in vitro* and *in vivo* screens, leading to a relative 2-fold increase in neurogenesis in old mice *in vivo*. Consistently, old NSCs exhibit a shift in glucose uptake, with 2 times more glucose uptake than their young counterparts. Reversing this shift, either genetically or by glucose starvation, boosted the ability of old NSCs to activate. Other stem cells, including hematopoietic stem cells and muscle stem cells, also display greater glucose usage and higher levels of glycolysis with age in mice and humans^90–92^, though the functional consequences of these glucose metabolism changes have not yet been tested. These observations suggest that increased glucose in cells may be a general feature of stem cell aging. This is interesting in light of the fact that glucose restriction extends lifespan in yeast^93, 94^ and worms^95, 96^, whereas high glucose shortens worm lifespan^97–99^. Importantly, dysregulated insulin-glucose signaling has also been linked to neuronal dysfunction and human brain aging and Alzheimer’s disease^100, 101^. Thus, our discovery that the increased glucose uptake in old cells can be targeted with GLUT4 knockout or glucose restriction raises the possibility that such mechanism could be targeted more broadly to counter aging.

Systemic gene therapy interventions can rejuvenate aspects of aging in progeria or even physiologically old mice^57, 102, 103^. A major challenge has been to rapidly identify new genetic interventions for rejuvenation *in vivo*. Our establishment of an *in vivo* screening platform shows a relatively high-throughput and rapid method of testing genetic interventions in the context of a regenerative stem cell niche *in vivo*. This type of screen should be scalable, versatile to use with other CRISPR-Cas9 techniques (e.g. gene activation/inhibition), and applicable to other cell types (e.g. other stem cells) in old mice. Genetic interventions that impact old tissues have the potential to identify new strategies – genetic or environmental – to delay or reverse features of aging. In the brain, such interventions can be particularly important to counter cognitive and regenerative decline during aging and neurodegenerative diseases.

## Acknowledgments

We thank L. Xu, J. Miklas, F. Boos, R. Nagvekar, S. Gagnon, and all Brunet lab members for their input on the project and providing feedback on the manuscript. We thank G.A. Reeves for independent code checking of the manuscript. Thank you to J. Butterfield and J. Ramirez-Matias for help with mouse husbandry and genotyping. We thank X. Ji for help with sequencing the *in vivo* screen libraries. This work was supported by P01AG036695 (A.B.), and a Larry L. Hillblom Foundation Postdoctoral Fellowship (T.J.R).

## Author contributions

T.J.R. designed the project with help from A.B., performed all experiments and computationally analyzed all results unless otherwise indicated, and wrote the manuscript with A.B. C.M.K. helped with NSC *in vitro* and *in vivo* immunofluorescence experiments, including virus production, mouse EdU injections, mouse perfusions, brain cryosectioning and processing for immunofluorescence and imaging. B.M. assisted with the *in vitro* genome wide screen experiments and assisted stereotaxic brain surgeries for *in vivo* screen experiments. R.W.Y. helped establish the lentiviral approaches for gene knockouts in NSCs, and trained C.M.K on mouse perfusions and cryosectioning. D.S.L. helped establish the qNSC culture reactivation model from young and old mice. D.W.M. provided feedback on the CasTLE analysis and helped design the *in vivo* screening experiments including the safe sgRNA controls. C.K.T. provided the plasmids and protocols for cloning sgRNAs. A.L. provided the plasmid libraries and technical input for genome-wide sgRNA lentivirus production. M.C.B. provided the sgRNA plasmid libraries, gave input on screen design, technical troubleshooting, and interpretation of screen results.

## Code availability

The code used to analyze screen data in the current study are available in the Github repository for this paper (https://github.com/Ruetz/Cas9_aging_NSC).

## Methods

### Laboratory animals

Cas9-expressing mice (Cas9 mice) were obtained from Jax (https://www.jax.org/strain/024858). These mice (background C57BL/6N) constitutively express the Cas9 endonuclease and an EGFP reporter under the control of a CAG promoter knocked into the *Rosa26* locus^1^. All screens in this study were performed with the Cas9 mice, including all NSC primary cultures and all *in vivo* work. We maintained a colony of Cas9 mice ranging in ages up to 28 months at the Stanford Comparative Medicine Building and the Neuroscience-ChemH building vivarium. As a negative control for the *in vivo* screens, male C57BL/6 mice obtained from the National Institute on Aging (NIA) Aged Rodent colony were used at 18-21 months old. NIA mice were habituated in the Stanford facility for at least 2 weeks prior to initiation of experiments. Mice were maintained under the care of the Veterinary Service Center at Stanford University under IACUC protocols 8661.

### Primary cultures of NSCs from young and old brains and reactivation experiments

For all experiments involving primary culture of NSCs, we pooled subventricular zones (SVZs) from pairs of male and female Cas9 mice, either 3-4 month-old mice (young) or 18-21 month-old mice (old). To generate primary cultures of NSCs from young and old mice, we micro-dissected SVZs into a small drop of PIPES buffer (pH 7.4), minced them in a 10cm tissue culture dish with ∼100 chops of a scalpel blade, and suspended the tissue in PIPES buffer prior to centrifugation for 5 minutes at 300g, at which point the excess PIPES buffer was poured out. The pellet of minced SVZs was then enzymatically dissociated (in 5 mL per 2 SVZs) with a mixture of HBSS (Corning, 21-021-CVR) with 1% penicillin-streptomycin-glutamine (Gibco, 10378-016), 1 U/mL Dispase II (STEMCELL Technologies, 07913), 2.5 U/mL Papain (Worthington Biochemical, LS003126), and 250 U/mL DNAse I (D4527, Sigma-Aldrich), vortexed briefly, and incubated at 37°C for 40 minutes on a rotator. The samples were then centrifuged at 300g for 5 minutes at room temperature and resuspended in NeuroBasal-A medium (Gibco, 10888-022) with 1% penicillin-streptomycin-glutamine (Gibco, 10378-016) and 2% B27 minus vitamin A (Gibco, 12587-010) and triturated ∼20 times, centrifuged and resuspended in complete ‘*aNSC media’* Neurobasal-A (Gibco, 10888-022) supplemented with 2% B27 minus vitamin A (Gibco, 12587-010), 1% penicillin–streptomycin–glutamine (Gibco, 10378-016), 20 ng/mL of EGF (Peprotech, AF-100-15), and 20 ng/mL of bFGF (Peprotech, 100-18B), placed in a humidified incubator, 37°C, 5% CO_2_. After 3-4 days, neurospheres emerged in the media and were passaged by dissociation with 1 mL Accutase (STEMCELL Technologies, 07920) for 5 minutes at 37°C, washed once with PBS, and resuspended in aNSC media. Neurosphere cultures were maintained with passaging every 2-3 days, and all experiments were performed in cultures of less than 10 passages. Details on passage numbers are provided in experimental sections below. For cultures of quiescent NSCs (qNSCs), the aNSC culture media was changed to remove EGF and add BMP4 (50 ng/mL)(Peprotech, 315-27). The Complete *‘qNSC media’* is: Neurobasal-A (Gibco, 10888-022) supplemented with 2% B27 minus vitamin A (Gibco, 12587-010), 1x penicillin–streptomycin– glutamine (Gibco, 10378-016), 50 ng/mL of BMP4 (Biolegend, 94073), and 20 ng/mL of bFGF (Peprotech, 100-18B). To induce quiescence, tissue culture plates were pre-treated with PBS (Fisher Scientific, #MT21040cv) containing 50 ng/mL Poly-D-Lysine (Sigma-Aldrich, P6407) for 1 hour, and then washed 3 times with PBS prior to cell plating cells on plates in qNSC media. The density of cells plated is important for induction of quiescence and ability of qNSCs to reactivate, especially in the context of lentiviral infection. In optimizing qNSC reactivation protocol, we observed that qNSCs seeded at the following densities were best for quiescence/reactivation experiments: 2×10^7^ cells per 15cm plate, 1×10^6^ cells per well of a 6-well plate, 2×10^5^ cells per well of a 24-well plate, and 1×10^5^ cells per well of a 96-well plate. For reactivation of qNSC cultures, cells were washed once with PBS, and then aNSC media was added to the plate, and refreshed every 2 days. For plating, cells were counted manually with a hemocytometer or using the Countess II FL Automated Cell Counter (Life Technologies, AMQAF1000).

### Tissue culture plastics

We found that primary cultures of NSCs were sensitive to the tissue culture plastic products used. Specifically, passaging NSCs in conical tubes manufactured by Genesee (15 mL conical tubes, Cat#28-103) resulted in death of the NSC cultures within 1 week of brief exposure to the plastic during passaging. Plastics from the following manufacturers were assessed to be suitable for NSC growth both in detached and adherent conditions: Thermo fisher 15/50 mL Falcon tubes (14-959-53A/14-432-22), 15cm/10cm /6-well/12-well/24-well/96-well Falcon® Tissue Culture Dishes (353025/08772E/08-772-1B/08-772-29/08-772-1/087722c).

### Lentivirus production

#### Genome-wide virus library preparation

For lentiviral production, human embryonic kidney 293T cells were seeded in DMEM + 10% fetal bovine serum (FBS, Gibco 10099141) + 1x penicillin-streptomycin-glutamine (Gibco, 10378-016) at a density of 2×10^7^ cells in 15cm plates. One day later, 293T media was replaced with 18 mL fresh media and the cells were transfected using the polyethylenimine (PEI) (1 mg/mL, Polysciences #23966-2) transfection method, mixing plasmids as follows: 2.27 μg each of 3^rd^ generation lentivirus packaging vectors pMDLg, pRSV, and pVSVG (obtained from Mike Bassik lab), along with 45 μg of the pooled single guide RNA (sgRNA) genome wide plasmid library (https://www.addgene.org/pooled-library/bassik-mouse-crispr-knockout/<u>). The sgRNA library targets all ∼23,000 protein coding genes in the genome, with 10 unique sgRNAs per gene, as well as 15,000 control sgRNAs (∼245,000 sgRNAs total)</u>^2^<u>. The sgRNA plasmid library, consisting of 20 sub-libraries, was mixed proportionally to the number of sgRNAs in each library. One day after PEI transfection, the medium was changed to 18 mL of NeuroBasal-A + 1x penicillin-streptomycin-glutamine (Gibco, 10378-016). After one day, the viral containing supernatant was collected on ice and stored at 4°C. Fresh medium was added to the 293T cells and collected again after 24 hours and again at 48 hours (a total of 3 collections of 18 mL of virus supernatant). All three supernatants were combined, filtered through a 0.45 μm filter (Stericup, EMD Millipore, #S2HVU02RE), and frozen at -80°C in 10 mL aliquots in 15 mL conical tubes. For plasmid library reamplifications, we electroporated 1 μL of 25 ng/μL of each library into 50 μL bacteria (Lucigen, 60242-2), with: 1.8 kV, 600 ohms, 10 μF in 0.1 cm cuvette (Gene Pulser Xcell, Bio Rad, 1652662). After electroporation, we allowed bacteria to recover in Lucigen recovery media for 2 hours in 15 mL conical tube shaking at 37°C. We plated 1 μL of the transformation onto an LB + carbenicillin (100 μg/mL, Sigma-Aldrich, #C9231-1G) agar plate to confirm transformation efficiency and the rest of recovery suspension was placed into 0.5 L LB + carbenicillin (100 μg/mL) liquid media in a 2L flask for 16 hr shaking at 37°C, and DNA was purified by Maxiprep (Thermo Fisher Scientific, FERK0492) according to the manufacturer’s protocol.</u>

#### sgRNA sub library design

We designed 5 sub libraries of sgRNAs to test gene hits from the *in vitro* screens in the brain (Fig. 1 and Fig. 2). Our selection criteria were as follows. For the **Top10 gene list**, we selected all significantly enriched genes (FDR<0.1) from the first 2 *in vitro* genome wide screens, selecting any gene that was significant in both screen 1 and 2, at any time point, day 4 or day 14 (for example, a Screen 1 day 4 hit and an overlapping Screen 2 day 14 hit would be added to the list). With that list, we ranked the genes based on the CasTLE gene score average from both screens and both time points (i.e. average of all: screen1 day 4 or day 14, screen 2 day 4 or day 14). This library was selected based on the first 2 *in vitro* genome-wide screens only, because at the time of library design, only the first 2 *in vitro* screens had been completed. Similarly, the **Depleted** gene list was also selected based on the first 2 screens. We selected all significantly depleted genes (FDR<0.1) from the first 2 *in vitro* genome-wide screens, selecting any gene that was significant in both screen 1 and 2, at any time point, day 4 or day 14 (for example, a Screen 1 day 4 hit and an overlapping Screen 2 day 14 hit would be added to the list). With that list, we then removed any gene that was significantly (FDR<0.1) depleted in qNSCs of screen 1 or 2 of any age. The final list was then selected by removing unannotated genes (e.g. GM3264, GM3164) and focusing on genes with associated publications. For the **Glucose uptake and Human disease** list, we selected genes that significantly (FDR<0.1) enriched in 2 of 3 *in vitro* genome-wide screens, at day 4 or day 14 (for example, a Screen 1 day 4 hit and an overlapping Screen 2 day 14 hit would be added to the list). The list of enriched genes was analyzed by GO term analysis (see section “Computational, analysis of CRISPR screens”). From the GO “Molecular Function (2018)” database, “D-glucose transmembrane transporter activity (GO:0055056)” and “sugar:proton symporter activity (GO:0005351)” terms were both in the top 10, with genes *Slc2a4, Slc2a12 and Slc45a4*. The other genes in the **Glucose uptake and Human disease** list were selected based on one of 2 criteria: (1) genes implicated in human disease: *Snrpb2* (Alzheimer’s)^3^, *Sorl1* (Alzheimer’s disease)^4^, *C1qtnf5* (Human aging)^5^, or (2) genes that are significantly (p<0.05) upregulated in qNSCs in old mice: *Slit2*^6, 7^*, Ier2*^7^*, Cdkn1a*^6, 7^*, Ecscr*^7^. For the **Cytoplasmic ribonucleoprotein granules** library, in the GO term analysis of gene knockouts that boosted old NSC activation, the terms “P-body (GO:0000932)”, “cytoplasmic ribonucleoprotein granule (GO:0036464)”, ribonucleoprotein granule (GO:0035770)”, “cytoplasmic stress granule (GO:0010494)” all came up in the list, although most were not significant. From this, we hypothesized that cytoplasmic granule structures could impede old NSC activation. We took the entire GO term “cytoplasmic ribonucleoprotein granule (GO:0036464)” and selected gene knockouts that had the greatest difference in effect between young and old NSC screens. Many of these genes did not demonstrate any significant effect in our *in vitro* screens, other than *Dis3l*2, *Edc3* and *Mbnl1*, which all significantly boosted old NSC activation in at least 2 of 3 screens. The final list of genes was the “**Published NSC regulators**” list, which we chose based on searching the literature for genes that had previously been implicated in regulation of NSC function and behavior. We did not select based on functional effect prediction.

#### sgRNA plasmid sub-library cloning for in vivo screens

The sgRNA expressing plasmid MCB320 (https://www.addgene.org/89359/) was digested with the BlpI and BstXI restriction enzymes, the band was gel-extracted and purified, and used for a pooled ligation reaction. We selected 5 sgRNAs from each gene of interest, based on the 5 out of 10 sgRNAs most enriched or depleted in our genome-wide *in vitro* screen. For the forward oligo of each sgRNA sequence, we added sequences: 5’-ttgg and 3’- gtttaagagc. For the reverse complement oligo of each sgRNA squence, we took the reverse complement of the sgRNA sequence and added 5’-ttagctcttaaac and 3’ -ccaacaag. To clone a pool of 10 genes, we selected the 50 sgRNAs pairs targeting the 10 genes and annealed the sgRNA pairs in separate annealing reactions, in a 100 μL of IDT duplex buffer (#11-05-01-12) with 1 µM forward and reverse oligos. We incubated the oligo pairs at 95°C for 5 minutes, and then allowed the oligos to gradually anneal at room temperature. We mixed all 50 annealed oligo pairs into one pool, diluted it 1:20 in IDT duplex buffer, and then used 1 μL of annealed oligo pool in a ligation reaction with 500 ng of digested MCB320 backbone. We used 1.5 μL of the ligation mix and electroporated 30 μL competent bacteria (Lucigen, 60242-2) with: 1.8 kV, 600 ohms, 10 μF in 0.1 cm cuvette (Gene Pulser Xcell, Bio Rad, 1652662). We plated the entire recovered transformed bacteria on a 10cm LB + Ampicillin (100 μg/mL, Sigma-Aldrich, #A9518-100G) plate, allowed overnight recovery, and the next day added 5 mL LB to the bacterial lawn and scraped it with a sterile silicon scraper. The resuspended bacterial mix was transferred to a clean collection tube, plate was again rinsed with additional 5 mL LB and transferred to same tube for overnight growth in 500 mL LB + Ampicillin (100 μg/mL) for Maxiprep (Thermo Fisher Scientific, FERK0492) according to manufacturer’s protocol. For library reamplification, we performed the same transformation and amplification procedure.

#### Concentration of virus for in vivo and in vitro sub-screens

For *in vivo* and *in vitro* sub-screens, we generated virus the same way as our genome-wide virus libraries, with modifications as follows. We plated 4 15cm plates of 293T cells for a total of 200 mL of collected virus after 3 days of collecting at 4°C, but rather than directly freezing the virus, we performed ultracentrifugation to concentrate the virus. For ultracentrifugation, we sterilized 30 mL ultraclear tubes (Beckman coulter 344058) under UV (TC room biosafety cabinet) for 15 minutes. We then put the tubes on ice, allowed 15 minutes to cool, and then added 30 mL of virus and centrifuged at 16,500 RPM for 1 hour at 4°C. We carefully decanted the supernatant using serological pipets, leaving 1 mL media in bottom of tube, adding 30 mL more virus-containing media and centrifuging again. We repeated the decanting, refilling and centrifugation of the same tube, concentrating a total of 180 mL of virus supernatant into a single tube. After the last ultracentrifugation, we removed most of the supernatant with a serological pipet, and the last 1 mL with a P1000 pipet tip from the side of the tilted tube, so as not to disturb the viral pellet. The viral pellet was usually visible in center of all of the tubes. We resuspend in 60 μL ice-cold PBS (1/3000^th^ original volume) by pipetting up and down ∼60 times, being careful not to produce air bubbles. The concentrated resuspended virus was then aliquoted into PCR strip tubes in 5 μL aliquots and placed onto dry ice. After 15 minutes, the virus was transferred to -80°C for storage. For experiments, virus was thawed on ice and injected into the brain or added to cell culture within 30 minutes of thaw. We assessed virus infectivity of each batch by performing serial dilution (3 μL, 1 μL, 0.5 μL) infections of 2×10^5^ 293T cells in 24-well culture plates for 16 hour infection, and then performing fluorescence-activated cell sorting (FACS) analysis 48-hours later to detect the percent of cells expressing mCherry reporter. For each experiment we normalized virus infectivity (viral titer) across treatments by adjusting the concentrations of virus added in PBS.

### Genome-wide knockout screens in primary cultures of NSCs

For each genome-wide screen, primary cultures of NSCs derived from a pool of 3 male and 3 female Cas9 mice were used for each independent biological replicate. In total, 3 independent genome-wide screens, each performed with independent young and old NSC pooled from 6 mice, were conducted. For each independent screen, young and old NSC cultures were processed in parallel at each stage of sample processing. The young and old NSC culture passage numbers were kept the same and at the start of screen were as follows: Screen 1, passage 8; Screen 2, passage 7; Screen 3, passage 12. To expand the NSCs up to the 1.4×10^9^ cells (the equivalent of 140 15cm plates) required for each biological replicate, 1×10^7^ NSCs were passaged and expanded into 15cm plates every 2-3 days, with feedings every 2 days (alternating between doubling the media (with 2x growth factor aNSC media) or complete media exchange). For each screen, 70 plates of 2×10^7^ qNSCs were seeded at day 0 (see below for library coverage calculations). After 4 days in quiescence media, the cells were incubated with the genome-wide sgRNA lentivirus library (see above). For this, sgRNA lentivirus library was freshly thawed at room temperature and diluted 1:5 in Neurobasal media and then B27 and growth factors were added to make it qNSC media, and 18 mL of this mix was added to plates for 16-hour overnight infection. The virus dilution added was based on viral titering experiments determined to achieve ∼30% culture infection of cells to ensure each cell only received a single sgRNA. Therefore, infecting the starting 1.4×10^9^ cells at 30% infection would result in 4.2×10^8^ infected cells, giving us a coverage of ∼1,700 cells per sgRNA (∼243,000 total sgRNAs). Note that these numbers represent the starting library coverage, but the cells do expand over the course of activation, and therefore the final cell numbers at end of experiment are orders of magnitude larger. The infected cells were then left in quiescence media for an additional 5 days prior to transition to aNSC media for activation. After 4 days of activation, the cells were dissociated using Accutase (Stem Cell Technologies, 07920) for 15-30 minutes at 37°C (until most cells rounded up) and gently scrapped with silicone cell scrapers (Fisher Scientific, 07-200-364) and split into 2 groups: 55 plates of NSCs were processed for the **Day 4** Ki67 FACS sorting (see below), and the other 15 plates of cells were placed into aNSC culture for 10 days of further expansion as neurospheres (**Day 14** time point). The day 4 FACS-sorted young and old cells were sorted to have equal numbers of Ki67^+^ cells from both ages for each screen for downstream analysis. See “Intracellular FACS” section below for Day 4 FACS protocol. The final number of sorted cells for each age in each screen was as follows: Screen 1 had 2.2×10^7^ sorted Ki67^+^ cells, Screen 2 had 1.41×10^7^ Ki67^+^ cells, Screen 3 had 1×10^8^ Ki67^+^ cells. After sorting, the methanol fixed cells were centrifuged at 700g for 5 minutes, the supernatant FACS buffer was decanted, and the cell pellets were frozen at -80°C until the genomic DNA extraction. To extract genomic DNA of sorted and methanol fixed cells, the cell pellets were defrosted at room temperature and then processed by resuspension in 5 mL of TE 1% SDS (Thermo Fisher Scientific, 15525017) and incubated at 65°C for 16 hours. The cell suspension was then treated with 50 μL proteinase K (Fisher Scientific, 25-530-049)(20 mg/mL) for 2 hours at 37°C. Samples were processed for genomic DNA extraction using Zymo Research ChIP DNA clean and concentrator (Zymo, D5205) according to manufacturer protocol. The day 14 expanding neurospheres were immediately centrifuged at 300g for 5 minutes and then processed for genomic DNA extraction with Qiagen QiaAmp DNA Blood Maxi Kit (51194), adding 5×10^7^ cells per column and according to the manufacturer’s protocol.

### sgRNA PCR amplification and sequencing

After genomic DNA isolation, sgRNA was amplified from the genome in 2 successive, nested PCR reactions. For the nested PCR reactions, we used either Herculase II Fusion Polymerase (Agilent, 600679) for screen 1, or Q5 DNA polymerase (Fisher Scientific, M0491L) for screens 2 and 3, and Q5 DNA polymerase for *in vivo* screens, according to manufacturer’s protocol. In optimizing this PCR reaction, we found that Herculase II Polymerase was outperformed by Q5 polymerase outperformed Herculase II Polymerase, Q5 polymerase requiring fewer PCR cycles to get more amplicon product, which is why we switched to Q5 DNA polymerase. We used 5 μg genomic DNA in 50 μL reactions to run on thermocycler. For the first PCR, we used primers MCB1562 (aggcttggatttctataacttcgtatagcatacattatac) and MCB1563 (acatgcatggcggtaatacggttatc) (1 µM final concentration), with PCR cycles as follows: 98^°^C for 2 minutes, 19 cycles of [98°C/30s, 59.1°C/30s, 72°C/45s], followed by 72°C for 3 minutes. We pooled all the PCR#1 cycle products and then used 5 µL of the pool in a second PCR reaction (PCR#2) with the same conditions but using different primers; MCB1439 (caagcagaagacggcatacgagatgcacaaaaggaaactcaccct) and a barcoded primer (aatgatacggcgaccaccgagatctacacGATCGGAAGAGCACACGTCTGAACTCCAGTCACXXXX XXCGACTCGGTGCCACTTTTTC, where XXXXXX is the 6-digit barcode for high throughput sequencing sample identification). The second PCR reaction was run for either 30 cycles (*in vitro* screen 1 and *in vivo* screens), or 18 cycles (*in vitro* screens 2 and 3). The resulting PCR products were all resolved on a 1.5% DNA agarose gel, the 272bp band was extracted (Qiaquick Gel extractions kit, #28706), eluted in 10 μL ultra-pure water (Invitrogen, 10977023), and assessed on a bioanalyzer (Agilent, Bioanalyzer 2100). Final libraries were combined into a pool at equal concentrations for sequencing on an Illumina Novaseq S4 system (with Novogene, for genome wide *in vitro* screen), or on an Illumina MiSeq system (with Stanford Genomics facility, for *in vivo* screens), sequencing to a depth of ∼1×10^7^ or ∼5×10^5^ reads per sample for *in vitro* and *in vivo* screens, respectively.

### Computational analysis of CRISPR/Cas9 screens

For both the *in vitro* and *in vivo* screens, our analyses were performed using CasTLE pipeline^8^. All of the source scripts can be found here: https://bitbucket.org/dmorgens/castle/downloads/. Briefly, for each screen, the raw screen fastq files were aligned to the sgRNA library sequence (“mm-Cas9-10”, or one of custom 10 gene library + control sgRNAs) to make count files using the “makeCounts” script. The count files were then analyzed using the “analyzeCounts” CasTLE script, comparing each screen timepoint to the starting plasmid sgRNA library count file (*in vitro* screens) or the sequenced 24-hour SVZ count file (*in vivo* screens), which we sequenced in parallel with screen libraries. We then calculated *P*-values for all genes in each screen by running 100,000 (*in vitro* screens) or 10,000 (*in vivo* screens) permutations with the “addPermutations” CasTLE script. For each genome-wide screen, we corrected for multiple hypotheses on the ∼23,000 gene associated *P*-values using the Python Statsmodel module, with Benjamini/Hochberg method, and classified genes as significant using an FDR<0.1 cutoff. For the *in vivo* screens, we classified genes as hits if their CasTLE computed 95% confidence interval did not contain 0. The library diversity of each sample was displayed using the “plotDist” CasTLE script. The screen results and individual gene sgRNA enrichment plots were visualized using the “plotVolcano” and “plotGene” scripts, respectively. To generate the final gene lists, we used all genes that were significant (FDR<0.1) in 2 or more independent screens (Screen 1 and 2, 2 and 3, or 1 and 3), at any time point (day 4 or day 14. For example-a Screen 1 day 4 hit and an overlapping Screen 2 day 14 hit, would be added to the list). For principal component analysis (PCA) we used the Python sklearn.decmposition.PCA module with CasTLE computed gene scores as input. We performed gene set enrichment analysis by inputting gene lists into the EnrichR online portal (https://maayanlab.cloud/Enrichr/) ^9, 10^, and then focusing on the “Ontologies” tab with GO Biological Process(2018)/Molecular function(2018)/Cellular(2018) components, sorting the terms based on *P*-value, which is computed by EnrichR using the Fisher exact test.

### Intracellular fluorescence activated cell sorting (FACS)

For the genome-wide screen and for other qNSC reactivation experiments, we FACS-isolated proliferative cells (Ki67^+^) as follows. Cells were dissociated with Accutase (Stemcell Technologies, 07920) for 5 minutes, collected into conical tubes, and centrifuged at 300g for 5 minutes. Cells were resuspended in PBS at 5×10^7^ cells in 1 mL (or 1×10^5^ cell in 100 μL), and then 9 mL (or 900 μL) ice-cold 100% methanol was added and cells were agitated for 15 minutes at 4°C. Cells were then centrifuged at 500g for 5 minutes and resuspended for a wash in 3 mL PBS and centrifuged again at 500g for 5 minutes. Cells were then resuspended in 3.5 mL staining solution: Ki67-APC (eBioscience, 17-5698-82) 1:300 in PBS, 2% fetal bovine serum (FBS) (Gibco, 10099141) at 4°C. Samples were agitated for 30 minutes at room temperature in the dark, and then 10 mL PBS was added prior to centrifugation at 700g for 5 minutes. Samples were then resuspended (25 mL per 5×10^7^ cells) in FACS buffer: PBS, 2% FBS, DAPI (Fisher Scientific, 62248, 1 mg/mL) 1:5000. Each sample was filtered with FACS-strainer cap tubes (Fisher, 08-771-23), just prior to FACS sorting. Cells were sorted on an Aria BD FACS Aria with a 100 μm nozzle at 13 psi and Flowjo (v10) software was used for data analysis.

### *In vitro* sub-screens and single gene knockout qNSC reactivation experiments

For testing the Top 10 gene library and single gene knockouts, we performed qNSC reactivation experiments in 24-well or 96-well format. We seeded 2×10^5^ cells in 24 well, or 1×10^5^ cells in 96 well format. After 4 days in quiescence media, with qNSC media changes every 2 days, concentrated virus was then added to the cells. We added 3 μL equal titer virus (see “Concentration of virus for in vivo and in vitro sub screens” section) to each 24-well containing 500 μL qNSC media, or 0.1 μL virus to 100 μL in each 96-well experiment. We left the virus in media with cells for 16 hours, and then refreshed the media. 5-6 days after infection, the cells were washed 1x in PBS and then transitioned to aNSC media for activation. aNSC media was exchanged once after 48 hours and then Ki67 intracellular FACS was performed at day 4 post infection (see intracellular FACS section above).

### *In vivo* gene knockout experiments

Stereotaxic surgeries were performed to inject virus into the lateral ventricle of mice. For these experiments, old Cas9 mice were used, except for one experiment where old WT mice were used (see “Laboratory animals” section). Surgeries were performed on heating pads with isoflurane induced anesthesia, with a Kopf (Model 940) stereotaxic frame, World Precision Instruments (UMP3T-1) UltraMicroPump3, Hamilton 1710RN 100 μL syringe with 30g Small Hub RN needle with point 2 beveled end. Injections were made at the following coordinates, relative to bregma: lateral 1 mm, anterior 0.3 mm, ventral depth 3 mm from skull surface. After drilling skull and inserting the needle into position, we waited 5 minutes prior to injecting virus. We injected 3 µL of equal titer virus at a rate of 10 nL/s. We waited 7 minutes after injection, before removing the needle and suturing the skin. Animals were administered a single dose of Buprenorphine SR (0.5 mg/kg) for postoperative pain management and monitored for 1-week post-surgery until full recovery. For labelling of proliferating NSC progeny, we injected animals intraperitoneally weekly with EdU (Thermo Fisher Scientific, A10044, 50 mg/kg, dissolved in sterile PBS), starting 1 week after surgery. We used both male and female mice for *in vivo* testing, always making note of sex for each experiment. We did not observe major differences in results between sexes, and plots include data from both sexes.

#### Influence of the anesthetic

In our pilot experiments, we performed some surgeries with Ketamine/Xylazine anesthesia instead of Isofluorane, for the relative ease of use, which we believe resulted in striking impairment of neurogenesis in both young and old animals when assessed in downstream screen analyses. Briefly, we performed our *in vivo* screening as outlined above, but we could only detect very few sgRNAs in the olfactory 5 weeks after injection when the mice had been anesthetized with Ketamine/Xylazine. We interpreted the lack of sgRNA detection in the olfactory bulb as an indication that not many NSCs were able to activate and migrate to the ofactory bulb in those conditions. We repeated the experiments with Ketamine/Xylazine 2 times, in close to 20 animals, always observing an impairment in sgRNA detection in the olfactory bulb after 5 weeks. When the same virus was injected into same age/background mice under isoflurane anesthesia, we could detect a far greater diversity and abundance of sgRNAs in the olfactory bulb 5 weeks later. We therefore did not perform any surgeries presented in this manuscript with Ketamine/Xylazine anesthesia, but used Isoflurane instead.

At the end point of *in vivo* experiments, mice were either sacrificed for sequencing of sgRNAs in the brain (*in vivo* sub-screens), or for immunofluorescence imaging (see *In vivo immunofluorescence experiments* section) of the olfactory bulb and other brain regions (single gene knockout experiments). For sequencing sgRNAs in the brain, mice were sacrificed either 1-2 days after injection or 5 weeks after injection and their brains were immediately removed and sub-dissected for genomic DNA extraction. We used a scalpel to cut off the olfactory bulbs and to cut a ∼1 mm thin slice of the outer cortex as well as the outer cerebellum. We then sub-dissected out the SVZ niche. We took each tissue and minced it with ∼100 cuts of a scalpel and proceeded to extract genomic DNA according to manufacturer’s protocol (Qiagen QIAamp DNA micro kit, 56304). The genomic DNA was then processed for sgRNA amplification and sequencing as outline in the *sgRNA PCR amplification and sequencing section*.

### Immunofluorescence staining of brain sections

For immunofluorescence straining of brain sections, young and old anesthetized mice were first subjected to intracardiac perfusion with 4 mL of heparin (Sigma Aldrich, H3149-50KU) and then 25 mL 4% paraformaldehyde (PFA) (Electron Microscopy Science, 15714) in PBS. Brains were then removed and further fixed for 16 hours by submerging in 4% PFA, at 4°C. Brains were then washed 3 times in PBS and placed in a conical tube with a 30% sucrose (Sigma-Aldrich, S3929-1KG) in PBS solution for 2-3 days until sinking to bottom of conical tube. The brains were then embedded in optimal cutting temperature (O.C.T.) compound (Electron Microscopy Sciences, 62550-12) for cryo-sectioning. Brain coronal sections were taken at 20 µm thickness (Leica, CM3050S). For assessing neurogenesis in the olfactory bulb, sections were taken every 200 µm across the entire olfactory bulbs. For SVZ imaging, we began taking slices at the most anterior part of the lateral ventricle, setting SVZ sections every 200 µm onto the slide. For immunofluorescence staining, sections were brought to room temperature and then washed 1 time with PBS and then permeabilized with ice-cold methanol and 0.1% Triton X-100 (Fisher Scientific, BP151) for 15 minutes. Slides were washed 3 times with PBS, and then treated with ClickIt reagents (for EdU) or put straight into antibody blocking solution. For Click-It EdU staining (Thermo Fisher Scientific, C10337/C10639/C10634), we placed 50-70 µL of reaction cocktail from this kit onto the tissue and incubated in humidified chamber at room temperature for 30 minutes. Slides were then washed 3 times in PBS prior to blocking for antibodies. Slides were treated with 50-70 µL blocking solution (5% normal donkey serum [NDS, ImmunoReagents, SP-072-V×10], 1% Bovine Serum Albumin (BSA, Sigma-Aldrich, A1595-50ML), 8.5 mL PBS) in a humidified chamber at room temperature for 30 minutes. Blocking solution was replaced with antibody solution consisting of blocking solution with antibodies as follows: mCherry (Invitrogen, M11217) 1:500, GFAP (Abcam, 53554) 1:500, GFP (Abcam, 13970) 1:500, GLUT4 (for *in vivo* staining R&D, MAB1262) 1:500, Ki67 (Invitrogen, 14-5698-082) 1:500, STX4A (Santa Cruz Biotechnology, sc-101301) 1:500, GFP (Abcam, 13970) 1:500, mouse IgG (Santa Cruz SC-3877, Lot: L1916) 1:500. After primary staining in dark for 2 hours in humidified chamber at room temperature or 16 hours at 4°C, slides were washed 3 times in PBS prior to staining with secondary antibodies. Secondary antibodies were diluted in blocking solution and consisted of Alexa 488/594/647 conjugated antibodies, (Fisher Scientific, A21202, A21206, A21209, A21447, A31571, A31573) 1:500, and DAPI (1 mg/mL, Fisher Scientific 62248) 1:5000. We added 50-70 µL of secondary antibody mix to cover the section, and incubated in dark for 2 hours in humidified chamber at room temperature or 16 hours at 4°C. Slides were then washed 3 times with PBS 0.2% tween for 10 minutes, washed 3 times with PBS for 5 minutes, and then mounted using ProLong Gold (20-40 µL, Thermo fisher Scientific, P36931), dried for 2 hours and sealed with nail polish. Images were captured using a Zeiss LSM 900 confocal microscope with a 10/20/63X objective. The exposure and gain settings for each channel/antibody were set at the beginning of each imaging session and remained the same for all animals and treatments. We randomized the order in which we imaged the slides, and we ensured that different treatments and age groups were all imaged in the same session on the same day. The imaging was not performed in a blinded manner. We did not select areas to image. We imaged and quantified the entire olfactory bulb or SVZ region (see “Immunofluorescence image analysis “section). For image analysis, see *Immunofluorescence image analysis* section below.

### Immunofluorescence staining of primary cell cultures of NSCs

For immunofluorescence staining of primary cell cultures of NSCs, we seeded 2.5 x 10^5^ aNSCs or 2×10^5^ qNSCs onto Poly-D-Lysine (50 ng/mL, Sigma-Aldrich, P6407) pre-treated (30 minutes, followed by 3x PBS wash) coverslips in each well of a 24-well plate. The qNSCs were plated 7 days prior to fixation, the aNSCs were plated 24 hours prior to fixation. For fixation, cells were washed 1 time with PBS, and then 500 μL of 4% PFA (Electron Microscopy Science, 15714) was added for 30-minutes incubation at room temperature. Cells were washed 3 times with PBS and then permeabilized with 0.1% Triton X-100 (Fisher Scientific, BP151)in PBS for 15 minutes shaking at room temperature. Coverslips were washed twice with PBS and then processed for antibody staining. Coverslips were placed on a 45 µL drop of primary antibody solution consisting of 1% BSA in PBS with primary antibodies as follows: GLUT4 (Abcam, 33780) 1:500, Ki67 (Invitrogen, 14-5698-082) 1:500, STX4A (Santa Cruz Biotechnology, QQ-17) 1:500. After 1-hour incubation in dark at room temperature, slides were washed 3 times in PBS shaking for 5 minutes at room temperature. Slides were then placed on 45 µL drop of secondary antibodies in 1% BSA in PBS consisting of Alexa 488/594/647 conjugated antibodies, (Fisher Scientific, A21206, A21209, A31571) 1:500, and DAPI (1 mg/mL, Fisher Scientific 62248) 1:5000. After 1-hour incubation at room temperature in the dark, slides were washed 3 times with PBS, prior to mounting with ProLong Gold, dried for 2 hours and sealed with nail polish. Images were captured using a Zeiss LSM 900 confocal microscope with a 10/20/63X objective. The exposure and gain settings for each channel/antibody were set at the beginning of each imaging session and remained the same for all samples and treatments. We randomized the order in which we imaged the slides, and we ensured that different treatments and age groups were all imaged in the same session on the same day. The imaging was not done in a blinded manner. We did not select areas to image. We randomly selected 10 areas of each coverslip to image. For image analysis, see *Immunofluorescence image analysis* section below.

### Immunofluorescence image analysis

For image analysis (both *in vivo* and in culture), we used the open-source software QuPath (https://qupath.github.io/) ^11^. For quantification of newborn neurons in the olfactory bulb, we first annotated a polygon line just beneath the olfactory bulb mitral cell layer, to focus the analysis within the inner layers of the olfactory bulb, where newborn neurons arrive. We then performed the “analyze cell detection” function, detecting cells in the image based on DAPI staining, using the program default settings, expanding the cell nuclei 5 µm in the “cell parameters” section. We then trained two independent object classifications: one for mCherry^+^ cells and the other for EdU^+^ cells, adjusting the thresholds to detect positive cells that were apparent by eye. We combined the mCherry and EdU objects into a single composite classifier and run it on all annotated images and treatments. The results were output as annotation measurements and annotation detections. The annotation measurements were used for graphs depicting the number of mCherry^+^EdU^+^/Total EdU^+^ cell numbers for each treatment, and the annotation detections were used to display the NeuN channel “cell mean” fluorescent intensity for EdU^+^mCherry^+^ populations in the different treatments. For quantification of GLUT4 fluorescent intensity *in vivo*, we first annotated a polygon line around the SVZ NSC niche, creating an analysis region about 5-20 cells deep from the ventricle wall. We then performed the “analyze cell detection” function, detecting cells in the image based on DAPI staining, using the program default settings, expanding the cell nuclei 5µm in the “cell parameters” section. We then trained two independent object classifications: one for Ki67^+^ cells and the other for GFAP^+^ cells, adjusting the thresholds to detect positive cells that were apparent by eye. We combined the Ki67 and GFAP objects into a single composite classifier and run it on all annotated images and treatments. The results were output as annotation detections. The annotation detections were used to display the GLUT4 channel “cell mean” fluorescent intensity for GFAP^+^Ki67^+^ (aNSCs), GFAP^+^Ki67^-^ (qNSCs/Astrocytes), GFAP^-^Ki67^+^ (Neuroblasts) and GFAP^-^Ki67^-^ (Other cells, including ependymal and microglia) populations across different aged mice. For the *in vitro* GLUT4 and STX4A quantifications, we selected the entire image as the analysis annotation. We then performed the “analyze cell detection” function, detecting cells in the image based on DAPI staining, using the program default settings, expanding the cell nuclei 5 µm in the “cell parameters” section. The results were output as annotation detections. The annotation detections were used to display the GLUT4 and STX4A “cell mean” fluorescent intensity for each protein’s channel in each cell-culture type and age group. For all experiments, the output numbers from different images were averaged across a biological replicate (1 mouse), biological sample values were then analyzed for significance by two-tailed Mann-Whitney test.

### Glucose uptake assays

For qNSC, we seeded 40,000 cells per well and for aNSC we seeded 10,000 cells per well (aNSCs do not stick to plate as well and will double every ∼16-24 hours, so we seed less to achieve similar density to qNSCs at time of analysis) on Poly-D-Lysine pre-coated 96-well plates, performing the assay 3 days after seeding. Duplicate wells were seeded and used for cell count normalization at time of glucose uptake assay. For knockout experiments, 1×10^5^ qNSCs were plated per well on Poly-D-Lysine (Sigma-Aldrich, P6407) pre-coated 96-well plates in quiescence media 6 days prior to infection with lentivirus to express sgRNA, where 1 μL of concentrated virus was added to the culture media for 16-hours to achieve ∼100% infection of the cells. We then assessed glucose uptake either 4 days or 8 days after infection. The glucose uptake assay was performed using either the 2NBDG (2-(*N*-(7-Nitrobenz-2-oxa-1,3-diazol-4-yl)Amino)-2-Deoxyglucose) (Fisher, N13195) fluorescent glucose analog (see below) or the Promega Glucose Uptake-Glo Assay (J1342) according to the manufacturer’s protocol, with the following details. Cells were pre-treated for 1 hour of qNSC/aNSC culture media without glucose. Culture media was then replaced with 50 µL of qNSC/aNSC media containing 1 mM 2DG (2-deoxy-D-Glucose, provided in Glucose Uptake-Glo kit from Promega (J1342)) reagent for 10 minutes in incubator (humidified, 37°C, 5% CO_2_). The 2DG media was then removed and 50 µL of PBS was added prior to carrying out the remainder of the assay according to the manufacturer’s protocol. All media treatments and reagent exchanges were pre-aliquoted into a empty 96 well plate, such that we could add the treatment to entire rows of cells at once using a multi-channel pipet, to ensure the duration of treatment was equivalent across different cell types and ages. The luminescence of the cells was measured with 0.5 second readings using a Varioskan LUX multimode plate reader. Due to different treatments having effects on cell numbers, plate readings in some cases (mentioned in figure legends) required normalization to the cell counts (Countess II cell counter, Thermo Fisher Scientific) based off duplicate wells. We performed glucose uptake experiments on different numbers of NSCs and observed a linear correlation between relative light units (RLU) and cells plated. For 2NBDG glucose uptake FACS assay, cells were placed in glucose free media for 1 hour and then treated with 200µM 2NBDG for 30 minutes at 37°C, and then analyzed by flow cytometry at excitation/emission maxima of ∼465/540 nm, with DAPI in the media to eliminate dead cells.

### Transient glucose starvation

NSCs were placed in qNSC media for 4 days, exposed to lentivirus to express sgRNAs targeting *Slc2a4* (GLUT4) or unannotated genomic regions (control). Then 6 days after infection, the cell media was replaced with standard complete qNSC media with glucose or modified to have no glucose (Neurobasal A media Thermo Scientific, A2477501, no D-glucose, no sodium pyruvate, supplemented with 1x sodium pyruvate, Fisher Scientific, 11-360-070) for 48 hours, at which point the media was replaced with standard complete aNSC media (with normal glucose concentration (4500 mg/L) in Neurobasal A media Thermo Fisher 10888-022) and the cells were allowed to activate for 4 days prior to intracellular FACS analysis with Ki67.

### Statistical analyses

For all experiments, young and old conditions were processed in an alternate manner rather than in two large groups, to minimize the group effect. We did not perform power analyses, though we did take into account previous experiments to determine the number of animals needed. To calculate statistical significance for experiments, all tests were two-sided Mann-Whitney tests. Results from individual experiments and all statistical analyses are included in Supplementary Table 4.

---

**Extended Data Figure 1.**
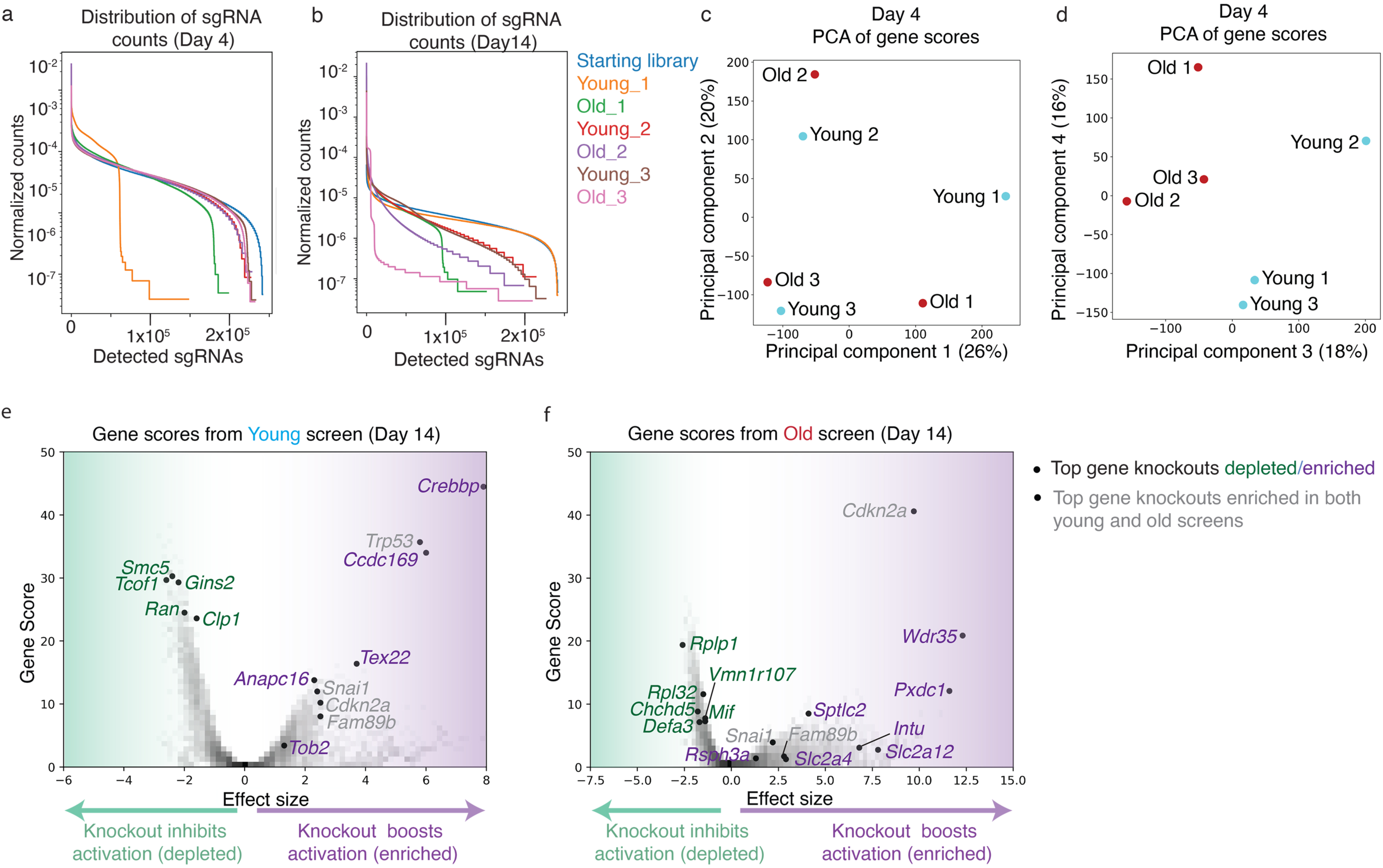
Genome-wide screen quality control. **a, b**, Normalized count matrices of all sgRNA counts across samples at Day 4 (**a**) or Day 14 (**b**). **c, d**, Principal Component Analysis (PCA) performed on all gene scores of the three independent screens at Day 4: Young 1, 2, 3 (blue) and Old 1, 2, 3 (red), with Principal Components 1 vs. 2 (**c**) and Principal Components 3 vs. 4 (**d**). **e, f,** Volcano plots of example screen results (screen 1) at Day 14 for young (**e**) or old NSCs (**f**), showing gene scores (y-axis) in relation to effect size (x-axis). Each dot represents one gene. Labelled dots are top ranking gene knockouts (FDR<0.1 in at least 2 of the 3 independent screens. Selecting genes that intersect screen 1 (day 4 or 14) with screen 2 (day 4 or 14)) that boost NSC activation (purple, corresponding to enriched sgRNAs) or impede NSC activation (green, corresponding to depleted sgRNAs) in an age-dependent manner or gene knockouts that boost activation regardless of age (grey, corresponding to enriched sgRNAs). See Supplementary Table 1 for complete list of gene scores.

**Extended Data Figure 2.**
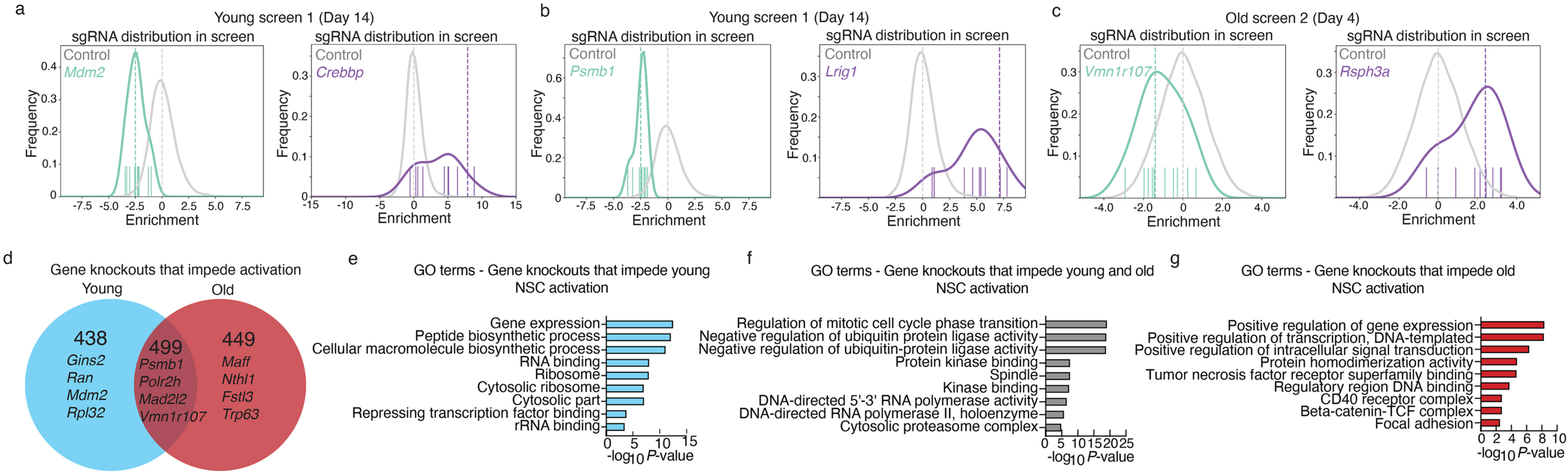
Example top sgRNAs counts from gene knockouts enriched or depleted in genome-wide screens and Gene Ontology analysis of gene knockouts that impede activation. **a-c,** Histograms plotting the relative enrichment and frequency of each sgRNA targeting a gene of interest or the control sgRNA pool (control), comparing the starting sgRNA plasmid library to the screen result. Enriched sgRNAs (purple) and depleted sgRNAs (green). Hashed line indicates the CasTLE computed enrichment effect size for sgRNA targeting the gene of interest and control sgRNA pool. Colored dashes above the x-axis represent each of the sgRNAs targeting the gene of interest and their relative enrichment. **d,** Venn diagram of all gene knockouts that impede NSC activation in at least 2 of the 3 independent screens (FDR<0. 1 in both screens. Selecting genes that intersect screen 1 (day 4 or 14) with screen 2 (day 4 or 14)) in young (blue) or old NSCs (red). **e-g,** Gene Ontology (GO) terms associated with gene knockouts that impede young (**e**) NSC activation, impede activation regardless of age (**f**), or impede activation of old (**g**) NSCs. For complete list of GO terms, see Supplementary Table 2. Gene sets selected based on FDR<0.1 in at least 2 of the 3 independent screens. GO terms assessed using EnrichR focusing on the “cell component”, “molecular function” and “biological process” libraries. *P-*value calculated by EnrichR using a Fisher’s exact test.

**Extended Data Figure 3.**
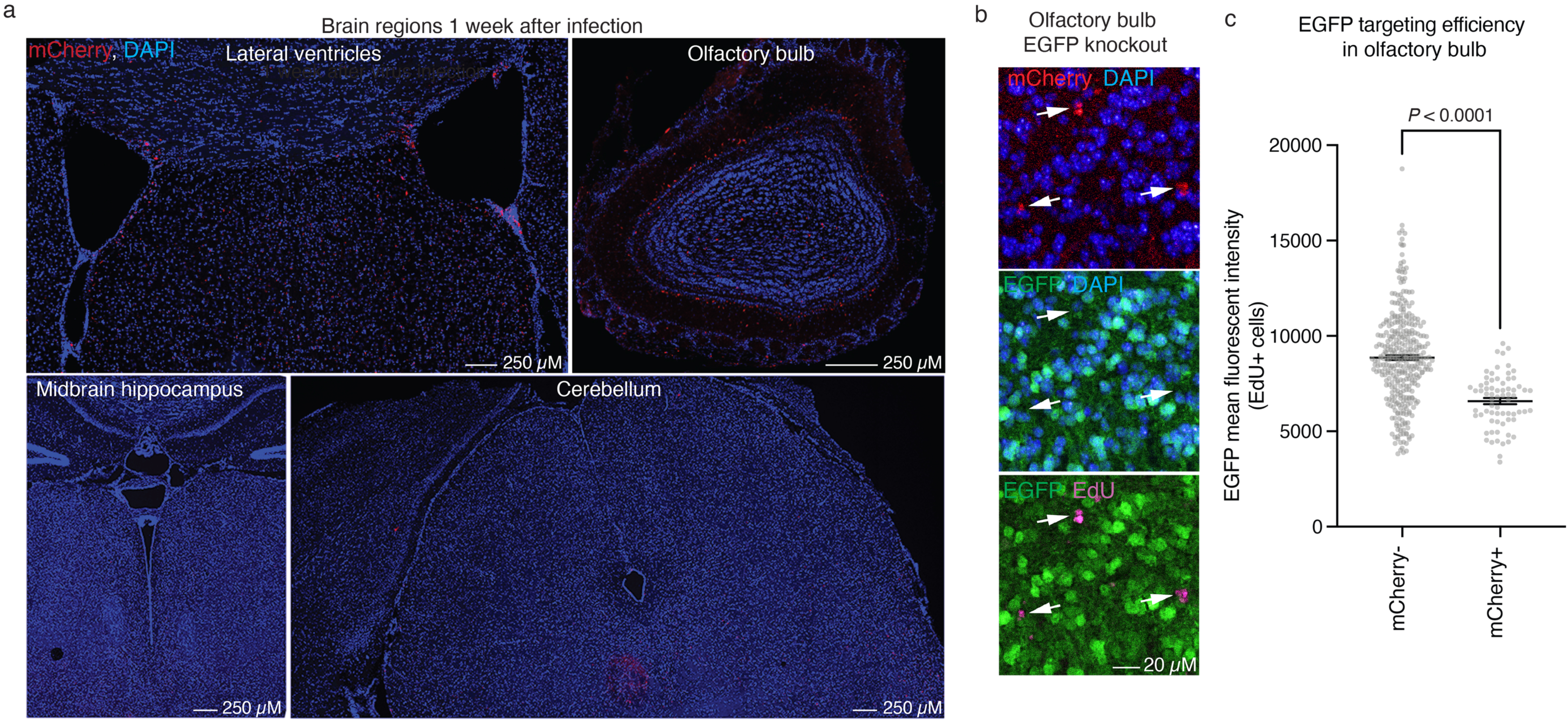
Efficiency of *in vivo* lentiviral injection and knockout. **a,** Immunofluorescence images of sections of the lateral ventricles and subventricular zones (SVZs), olfactory bulb, midbrain hippocampus and hindbrain cerebellum regions of old (22 months old) female Cas9 mice, 1 week after injection of lentivirus expressing mCherry reporter and sgRNAs. Markers: mCherry reporter (lentivirus-infected cells, red) and cell nuclei (DAPI, blue). **b**, Zoomed-in immunofluorescence images of sections of olfactory bulb from old (22 months old) female Cas9 mice, 5 weeks after injection of lentivirus expressing sgRNA targeting the EGFP reporter present in Cas9 mice directly into the lateral ventricles. Mice were injected with EdU once per week, starting one week after virus injection, for 4 weeks. Markers : mCherry reporter (lentivirus-infected cells, red), EGFP reporter (present in Cas9 mice, green), EdU (newborn cells, magenta) and cell nuclei (DAPI, blue). **c,** Quantification of EGFP mean fluorescent intensity in non-infected (mCherry^-^) and infected (mCherry^+^) newborn cells (EdU^+^) in the olfactory bulb, 5 weeks after injection of lentivirus to express sgRNA targeting EGFP into the lateral ventricle. Dot plot showing mean +/-SEM of the EGFP fluorescence intensity in EdU+ cells. Each dot represents a cells EGFP mean fluorescent intensity. *P*-value determined by two-tailed Mann-Whitney test.

**Extended Data Figure 4.**
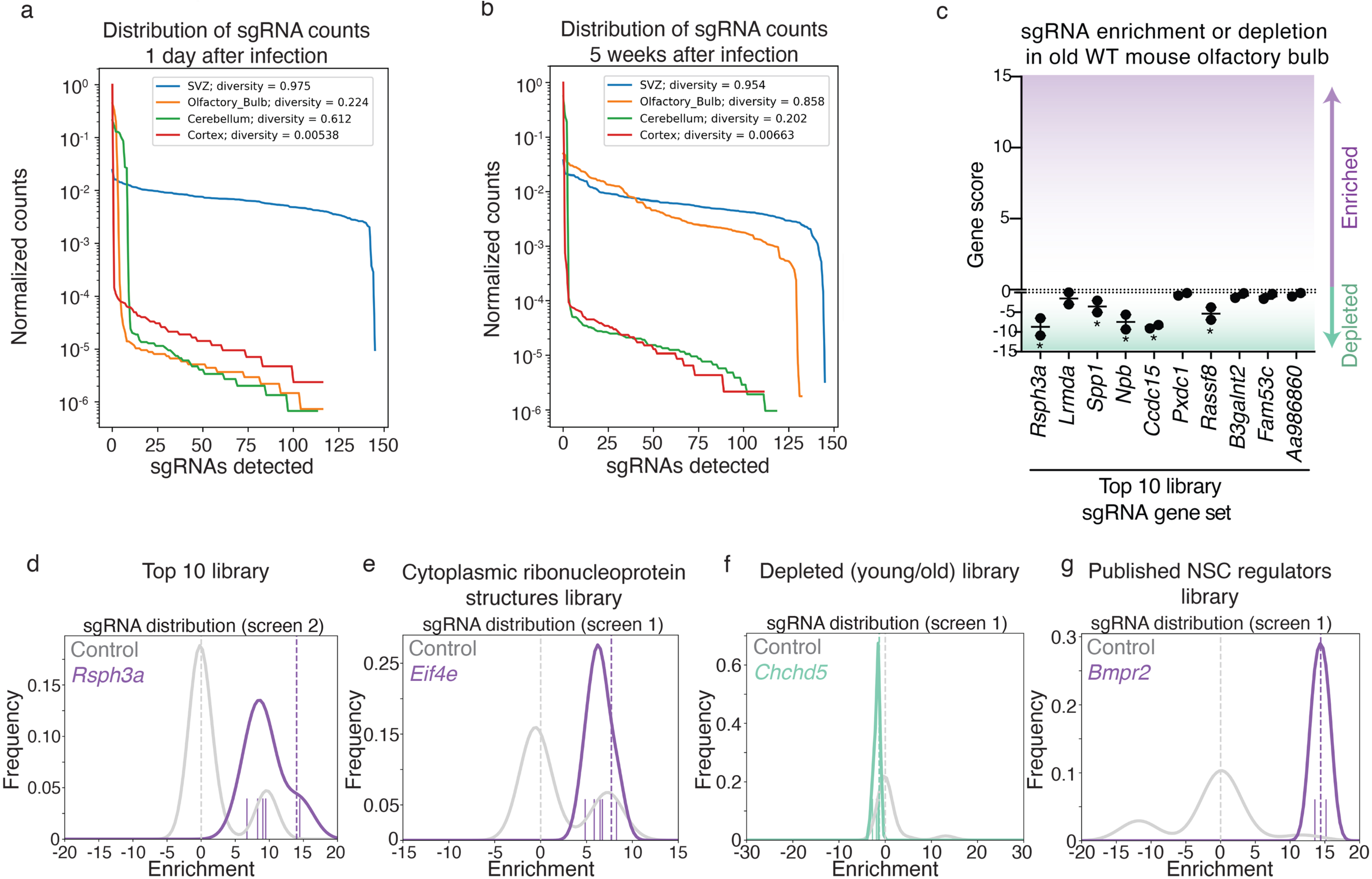
*In vivo* screen quality control, WT controls, and example genes. **a, b,** Normalized count matrices of all sgRNA counts across samples at 24-hours (**a**) or 5 weeks (**b**) post virus injection. **c,** Olfactory bulb sgRNA enrichment CasTLE analysis results showing gene scores of the Top 10 gene pool, 5 weeks after injection of lentivirus expressing sgRNAs targeted to these genes directly into the lateral ventricles of wild-type (WT) old (20-21 months old) male mice. Gene scores were computed by comparing the olfactory bulb sgRNA counts 5-week post injection and the SVZ sgRNA counts from an independent mouse sequenced 24 hours after injection. Each dot represents gene score from an independent mouse. * Denotes gene hits with a 95% confidence interval that did not contain 0 as computed by CasTLE analysis. **d**, Histograms plotting the relative enrichment and frequency of each sgRNA targeting a gene of interest or the control sgRNA pool, comparing the starting sgRNA library in the SVZ at 24 hours post infection to the olfactory bulb sgRNA counts at 5 weeks post injection. Here we show top ranked genes plotted on example screens as labelled. Hashed line indicates the CasTLE computed enrichment effect size for targeted gene and control pool. Colored dashes above the x-axis represent each of the sgRNAs targeting the gene of interest and their relative enrichment. See also Supplementary Table 3.

**Extended Data Figure 5.**
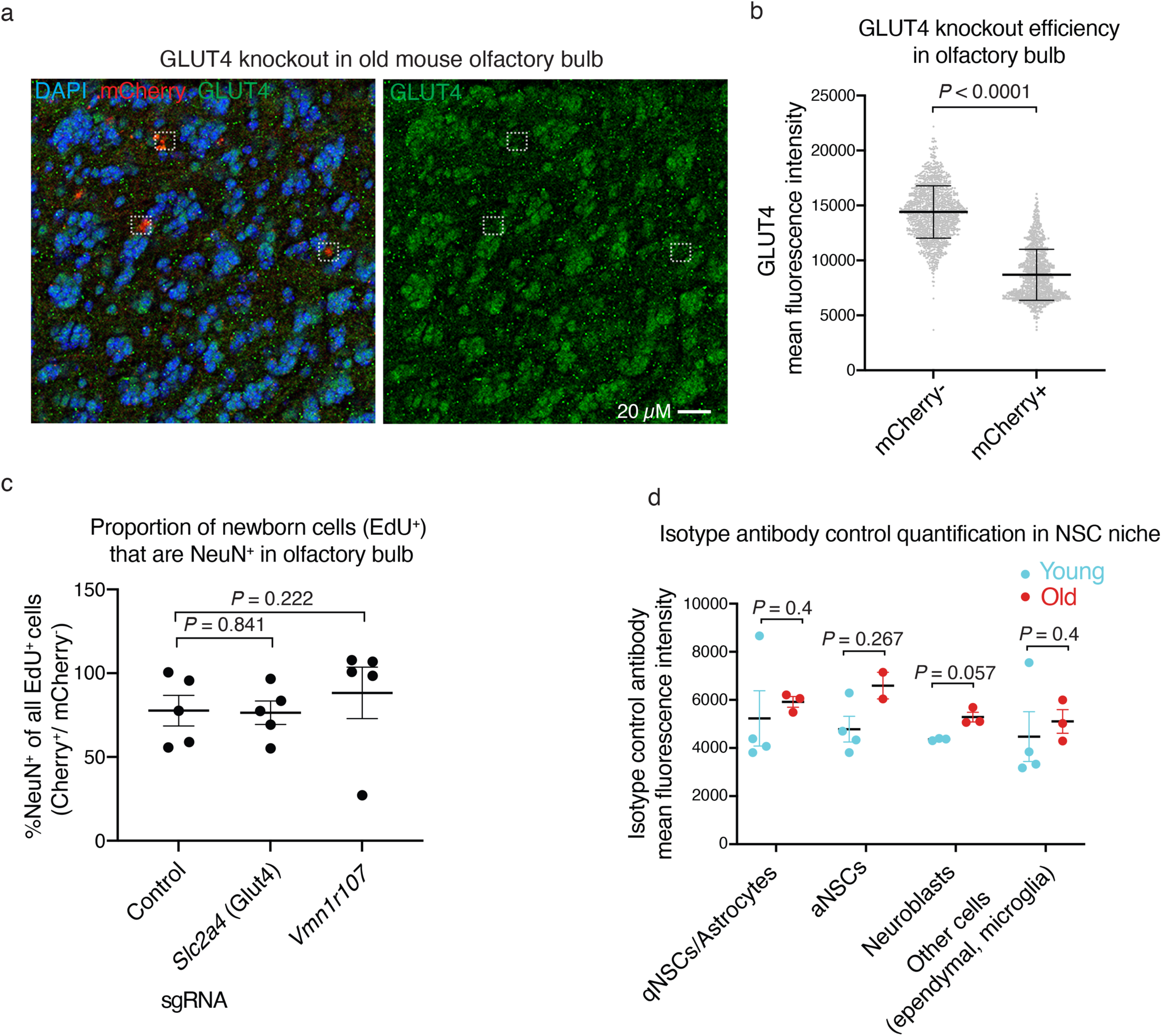
GLUT4 knockout efficiency *in vivo* and proportion of newborn cells that become neurons. **a,** Example immunofluorescence images of olfactory bulb sections from old (22 months old) male Cas9 mice, 5 weeks after injection with lentivirus expressing *Slc2a4* (GLUT4) sgRNAs directly into the lateral ventricles. Markers: mCherry (lentivirus-infected cells, red), GLUT4 (green), and DAPI (nuclei, blue). Dashed white squares are examples of cells with mCherry infection (expressing sgRNA targeting EGFP). **b,** QuPath image quantification of GLUT4 mean fluorescence, comparing infected (mCherry^+^, expressing sgRNA targeting GLUT4) and uninfected (mCherry^-^) cells. Dot plot showing mean +/- SEM of all detected cells GLUT4 mean fluorescence. Each dot represents a cell quantified in a *Slc2a4* (GLUT4) sgRNA treated mouse. *P*-value determined by two-tailed Mann-Whitney test. **c,** QuPath image quantification of the percentage of newborn cells (EdU^+^) that are also NeuN^+^, comparing cells with (mCherry+) or without (mCherry-), with *Slc2a4* or *Vmn1r107* sgRNA expression (see also Fig 3b). Dot plot showing mean +/- SEM of quantification results from 3 independent experiments, each with 1-3 mice, for a total of n = 5 mice.. Each dot represents the average (%NeuN^+^) of all cell quantifications from a single mouse. *P*-values determined by two-tailed Mann-Whitney test. **d,** QuPath image quantification of the mean fluorescence intensity with an isotype antibody control (mouse IgG) in sections from the SVZ neural stem cell niche from 4 young (3-4 months old) and 3 old (18-21 months old) male Cas9 mice (control for Fig. 3f). Cell types were identified as follows: qNSC/astrocyte (GFAP^+^/Ki67^-^), aNSC (GFAP^+^/Ki67^+^), Neuroblast (GFAP^-^/Ki67^+^) and other cells (ependymal, microglia; GFAP^-^/Ki67^-^). Dot plot showing mean +/- SEM of quantification results from 2 independent experiments, each with 1-3 mice, for a total of n = 3 or 4 mice. Each dot represents average fluorescent intensities of cells from one mouse. *P*-values determined by two tailed Mann-Whitney test.

**Extended Data Figure 6.**
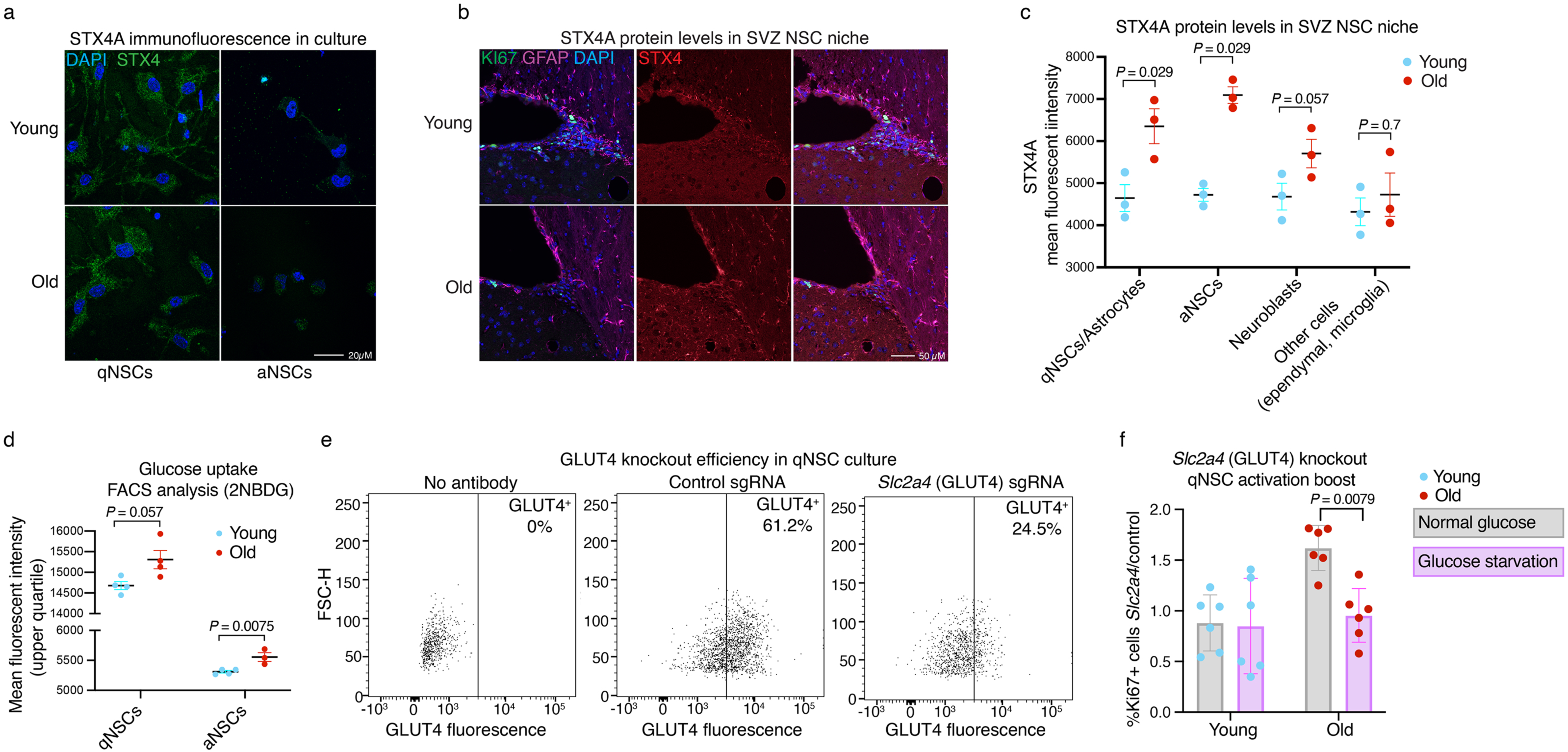
STX4A immunofluorescence *in vitro* and *in vivo*, *Slc2a4* (GLUT4) knockout efficiency and effects of *Slc2a4* (GLUT4) knockout on qNSC activation in the context of glucose restriction. **a,** Immunofluorescence image of STX4A in primary NSC cultures from young (3-4 months old) or old (18-21 months old) mice (quantification in Fig. 4e). NSCs were plated in quiescence NSC media (qNSCs) for 7 days prior to imaging, and NSCs were plated in activated NSC media (aNSCs) 2 days prior to imaging. Markers: STX4A (green) and DAPI (nuclei, blue). **b**, **c,** Representative immunofluorescence images of coronal sections from SVZ NSC niche sections from young (3-4 months old) and old (18-21 months old) Cas9 mice. Markers: Ki67 (proliferation maker, green), GFAP (NSC and astrocyte marker, magenta), DAPI (nuclei, blue), and STX4A (red). Cell types were identified as follows: qNSC/astrocyte (GFAP^+^/Ki67^-^), aNSC (GFAP^+^/Ki67^+^), Neuroblast (GFAP^-^/Ki67^+^), and other cells (ependymal, microglia; GFAP^-^/Ki67^-^). (**c**) QuPath image quantification of STX4A mean fluorescence in cells of the neural stem cell niche from 3 young (∼4 months-old) and 3 old (∼19 months-old) male Cas9 mice. Cell types were identified as follows: qNSC/astrocyte (GFAP^+^/Ki67^-^), aNSC (GFAP^+^/Ki67^+^), Neuroblast (GFAP^-^/Ki67^+^) and other cells (ependymal, microglia; GFAP^-^/Ki67). **d**, Glucose uptake assay with 2NBDG (2-(*N*-(7-Nitrobenz-2-oxa-1,3-diazol-4-yl)Amino)-2-Deoxyglucose) FACS on primary NSC cultures (qNSCs and aNSCs) from young (3-4-months-old) or old (18-21-months-old) mice. Dot plot showing mean +/- SEM of results from 3-4 independent cultures, each from a pool of 6 mice, 3 males and 3 females. Each dot represents an independent NSC culture. *P*-values determined by two-tailed Mann-Whitney test. **e**, GLUT4 knockout efficiency *in vitro*. FACS analysis with the GLUT4 antibody of qNSCs treated with control sgRNA (targeting unannotated regions of the genome) or sgRNA targeting *Slc2a4* (GLUT4), 10 days after lentivirus infection to express sgRNA. No antibody control panel is on the left. Plots show mCherry+ gated cells, GLUT4 fluorescence. **f**, Data from Fig. 4h, presented as the boost in qNSC activation ability with *Slc2a4* (GLUT4) knockout, with or without glucose starvation. Dot plot showing mean +/- SEM of activation ability of *Slc2a4* (GLUT4) knockout relative to control. Each dot represents an independent NSC culture. *P*-values determined by two-tailed Mann-Whitney test.

## Supplementary Tables

Supplementary Table 1: Gene scores for *in vitro* genome-wide screens

Supplementary Table 2: GO analyses of *in vitro* genome-wide screens

Supplementary Table 3: Gene scores for *in vivo* screens

Supplementary Table 4: Experimental data

